# Transcriptome-wide mapping reveals a non-canonical RNA-dependent mechanism of platinum cancer drugs

**DOI:** 10.64898/2025.12.20.694502

**Authors:** Arun Krishnaraj, Xinrui Wei, Richi Thakral, Weiyi Guo, Siva Bala Subramaniyan, Xiaopei Zhang, Katelyn R. Alley, Anqi Feng, Andrew Hoy, Douglas Kung, Purushottam B. Tiwari, Aykut Üren, Rodrigo Maillard, Victoria J DeRose, Sreejith J Nair

## Abstract

Off-target interactions frequently compromise the clinical utility of anticancer agents by driving dose-limiting toxicity and therapeutic resistance. Although RNA has been predicted to be an off-target for numerous FDA-approved drugs, the extent and functional significance of RNA off-targeting among anticancer small molecules remain poorly understood. Using a systematic drug-binding screen, we identified cisplatin, a frontline chemotherapeutic that acts canonically through DNA adduct formation, as a prominent RNA binder. We employed cisplatin as a model compound to characterize the mechanistic basis and functional impact of RNA-small molecule off-targeting. To map transcriptome-wide cisplatin-RNA interactions, we developed PlatRNA-seq, a click-chemistry-enabled RNA-binding profiling platform. Genomic and functional analyses reveal that cisplatin preferentially accumulates at RNA G-quadruplex (rG4) structures near 5′ transcript ends, inducing R-loop formation. Critically, we demonstrate that cisplatin cytotoxicity is partially mediated through RNA binding, revealing a noncanonical mechanism of action. Collectively, these findings illustrate the functional consequences of RNA-small-molecule off-targeting and provide a generalizable framework for investigating small-molecule-RNA interactions, opening new avenues for therapeutic innovation.

## Main

Despite significant progress in rational drug design and clinical research, several clinically used anticancer compounds exhibit limited efficacy and excessive toxicity ^1^. Off-target interactions with undesirable drug binders are a common cause of these side effects. Recent research shows that many small-molecule oncology drugs, initially believed to act through selective binding to single protein targets, often exert their therapeutic effects by engaging multiple targets ^2,3^. Adding to this complexity, although most anticancer small molecules are designed for proteins or DNA, emerging data suggest that a significant fraction are also predicted to bind RNA^4^. However, characterization of the structural basis and functional consequence of RNA off-targeting by any clinically used anticancer small molecule has not yet been undertaken. In parallel, there are growing efforts to target RNA structures with small molecules as novel therapeutic modalities for previously “undruggable” targets ^5–8^. Although predicting RNA-small molecule binding remains a major challenge, precise targeting of RNAs can increase the potential number of drug targets by orders of magnitude ^6^. Our work is motivated by the premise that systematically characterizing the biochemical and biological consequences of RNA interactions with common anticancer agents will reveal novel mechanisms of action, facilitating innovative therapeutic and prognostic opportunities in cancer treatment.

To experimentally determine the RNA off-target interactions of anticancer small molecules, we performed a targeted screen of a cohort of anticancer drugs. Under our assay conditions, approximately 9% of the compounds exhibited measurable RNA binding. Among these, cisplatin, a widely used chemotherapy agent for treating solid tumors, was consistently identified as an RNA binder. Cisplatin and its analogues, carboplatin and oxaliplatin, account for treatment in nearly half of all patients receiving chemotherapy worldwide ^9,10^. The cytotoxicity of cisplatin is classically attributed to the formation of DNA adducts, which trigger the DNA damage response (DDR) pathways and apoptosis ^11,12^. Biochemical evidence from lower eukaryotes and in vitro systems indicates that cisplatin also binds to structured, abundant, non-coding RNAs, such as tRNAs and rRNAs ^13–17^. However, there is no transcriptome-wide information on the sequence and structural determinants, or the functional consequences, of RNA interactions with cisplatin in mammalian systems. Thus, the extent and impact of such potential interactions and their relationships to cancer treatments remain largely unknown. Here, we show that RNA constituted a major off-target of cisplatin in human cancer cells. Using a novel chemico-transcriptomic assay (PlatRNA-seq), we report that cisplatin preferentially binds a subset of cellular RNAs at guanine-rich motifs that form noncanonical RNA G-quadruplex (rG4) structures prone to R-loop formation. Competitive inhibition of cisplatin recruitment to rG4s markedly reduces its cytotoxicity, indicating that RNA interactions contribute to its pharmacological activity. Collectively, these findings uncover a noncanonical RNA-mediated mechanism underlying the action of platinum-based chemotherapeutics, demonstrating that RNA off-targeting by anticancer small molecules carries direct functional consequences. More broadly, this work suggests that RNA off-targeting is a critical, yet overlooked mechanism of drug action and provides a framework for their systematic functional characterization.

### RNA off-target binding is prevalent among oncology drugs

To estimate the prevalence of RNA-binding specifically among oncology drugs, we assessed RNA-binding properties in the set of all FDA-approved oncology drugs (Fig. 1a, Supplementary Table 1), finding that ∼15% of oncology-related small molecules share structural similarity (Tanimoto coefficient, Tc ≥ 0.85) ^4,18^ with at least one validated RNA binder ^19,20^. To experimentally assess RNA interactions, we screened a library of 179 approved oncology drugs, of which ∼19% showed structural similarity to known RNA binders (Supplementary Fig. 1a). We utilized surface plasmon resonance (SPR) to screen this library for compounds that directly bind to RNA in real-time under label-free conditions (Fig. 1b). We tested two RNA libraries, one with random sequences and one with RNA homopolymer, against small molecules at concentrations ranging from 2 to 50 µM. Molecules showing between 25% and 200% of the expected Rmax were classified as RNA binders. Among the 179 compounds, 15 (8.3%) and 17 (9.5%) bound to the random and homopolymer RNA pools, respectively (Fig. 1c and Supplementary Fig. 1b–d). A total of 19 RNA-binding drugs were identified (10.6% of screened drugs), 13 of which were shared across RNA pools, demonstrating high reproducibility of the hits (Supplementary Fig. 1c–d). Two of the hits in our screen are listed in the combined RNA binder small molecule list (Supplementary Fig. 1d, highlighted in red). The experimental validation of RNA binders is less than predicted by the structural similarity estimate, which may reflect an overestimation from similarity calculations or the relative simplicity of the assay conditions. Taken together, these results demonstrate that RNA off-target interactions constitute a recurrent and underappreciated feature among approved oncology drugs.

**Fig. 1.**
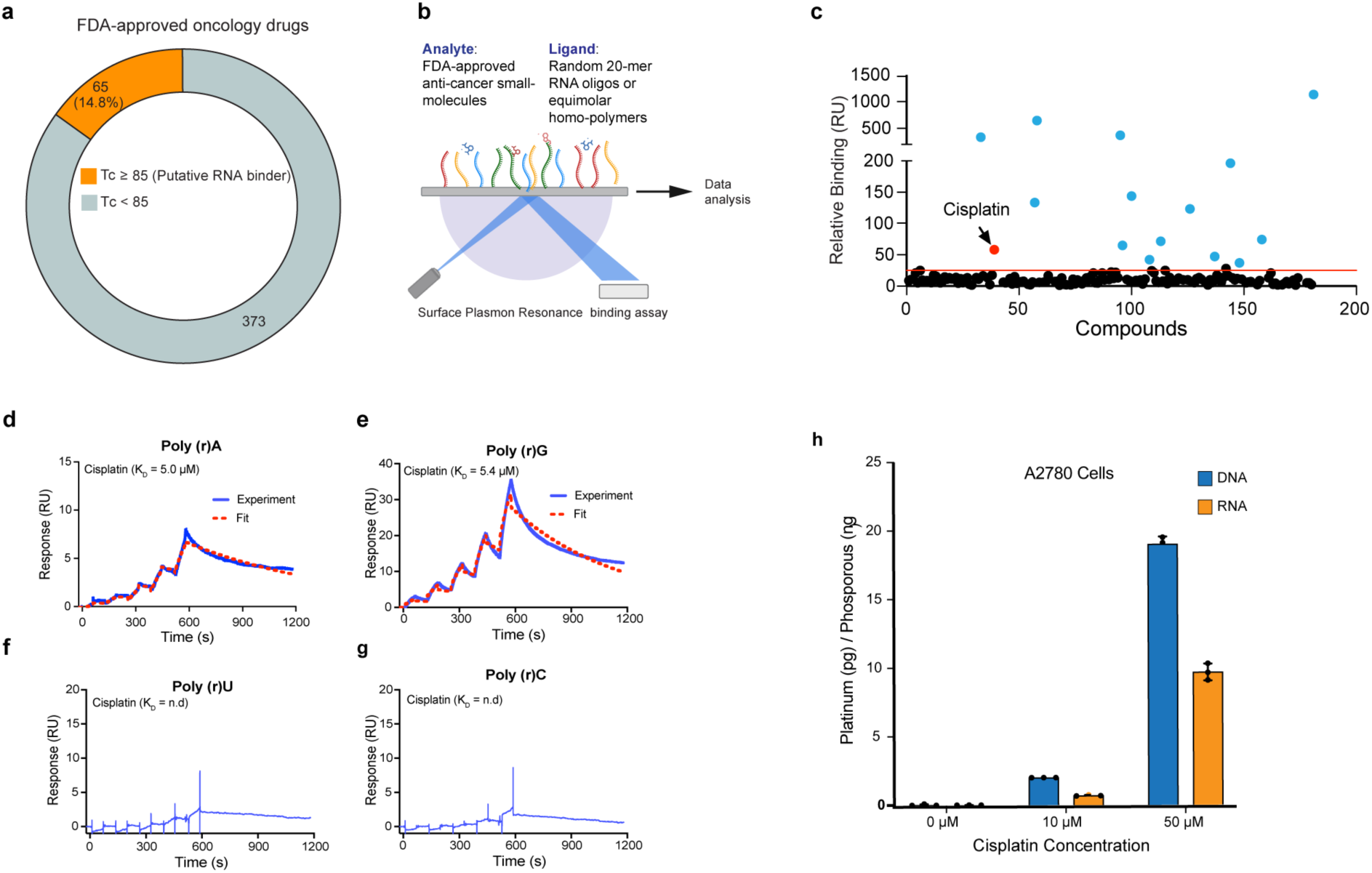
Cisplatin preferentially accumulates in cellular RNAs within cancer cells. (**a)** Distribution of FDA-approved anticancer small molecule drugs based on the structural similarity to at least one known RNA binder (Tanimoto coefficient, Tc >85). (**b)** Outline of the SPR-binding screen performed to identify RNA-binding of FDA-approved small-molecule oncology drugs. (**c**) The relative binding (Resonance Units) of individual small molecules in the library to the captured 20-mer RNA pool. The red line indicates the cut-off value to determine hits (please see the methods). (**d,e,f,g)** Single-cycle SPR data showing interaction between homopolymer RNA oligos and cisplatin. The binding affinity of cisplatin with each oligo is shown. n.d = not determined. Data representative of replicate experiments. (**h**) Quantitation of cisplatin accumulation in the DNA and RNA fractions of A2780 cell line.

### RNA is a major off-target of cisplatin in cancer cells

Among the identified hits, cisplatin was of particular interest. Cisplatin and its analogs remain among the most widely used chemotherapeutics globally ^21,22^. Yet, despite decades of clinical use, platinum-based therapies continue to face challenges such as poor tolerability and frequent emergence of resistance ^11,23^. Despite ongoing research, primarily focusing on the DNA-based action of cisplatin, these problems remain largely unresolved ^11^. Importantly, prior biochemical studies in lower organisms have shown that cisplatin can interact with abundant RNA species such as tRNAs and rRNAs ^13–15,17^, but the biochemical basis and functional relevance of these interactions in cancer patients remain unclear.

SPR kinetic assays using RNA homopolymers confirmed the preferential binding of cisplatin to purines ^15,24^ (Fig. 1d–g). The measured binding affinities (3-7 μM) fall within clinically relevant plasma concentrations observed in platinum-treated patients (2–50 μM) ^25–27^. To quantify cisplatin distribution between RNA and DNA in cells, ovarian cancer lines A2780 and TOV112D were treated with cisplatin, followed by isolation of DNA and RNA after removing contaminating nucleic acids, and subjected to inductively coupled plasma mass spectrometry (ICP-MS) analysis ^28^ (Supplementary Fig. 1e). In both cell lines tested, platinum accumulation increased in both DNA and RNA fractions in a dose-dependent manner (Fig. 1h, and Supplementary Fig. 1f). Although cisplatin accumulated approximately twofold more per nucleotide in DNA than RNA, a typical mammalian cell is estimated to contain 6-10-fold more RNA than DNA ^29,30^. Extrapolating from the per-nucleotide accumulation of cisplatin and the relative RNA: DNA abundance indicates that, in cancers treated with cisplatin, the majority of nucleic acid-bound drug (over 80%) may accumulate in RNA rather than DNA (Supplementary Fig. 1g). These results, together with the previous report ^13,31^, demonstrate that RNA is a major intracellular target of cisplatin in cancer cells.

### Development of PlatRNA-seq

While multiple approaches exist to map genome-wide DNA-platinum interactions, including HMGB1 domain A-based pulldown of cisplatin-linked DNA ^32^, antibody-based enrichment of cisplatin adducts ^33^, and indirect detection via nucleotide excision repair assays ^34^, no method has been developed to comprehensively map cisplatin interactions across the RNA transcriptome. Transcript-level detection of platinum-RNA adducts has been limited to targeted reverse transcriptase (RT) primer extension assays mainly on highly abundant ribosomal and tRNAs in lower organisms ^17,35–37^. By contrast, the unbiased evaluation of platinum adducts on regulatory RNAs such as mRNA and lncRNAs is more challenging due to several factors including lower abundance and heterogeneous accessibility. Furthermore, classical approaches using modified biotinylated platinum compounds for enrichment ^38,39^ can interfere with the coordination chemistry and reactivity of these small, non-carbon-based drugs ^40^.

To address these limitations, we developed a click-chemistry-based transcriptome screening approach. Click chemistry enables minimally disruptive small-molecule functionalization, thereby avoiding the steric effects associated with bulkier tags. The azide group is compact, bioorthogonal, and easily introduced via nucleophilic substitution without compromising drug activity ^41^. For RNA applications, copper-free strain-promoted azide–alkyne cycloaddition (SPAAC) is available to avoid copper-induced degradation. We therefore synthesized a clickable cisplatin analog, 1,3-platin, to label RNA targets of platinum drugs *in vivo* (Fig. 2a and Supplementary Fig. 2a), and validated concentration-dependent in-cell RNA binding and click efficiency on extracted RNA (Supplementary Fig. 2b–c). These results demonstrate that 1,3-platin interacts with cellular RNAs and, when paired with click labeling, enables the detection of RNA-bound platinum in its native environment.

**Fig. 2.**
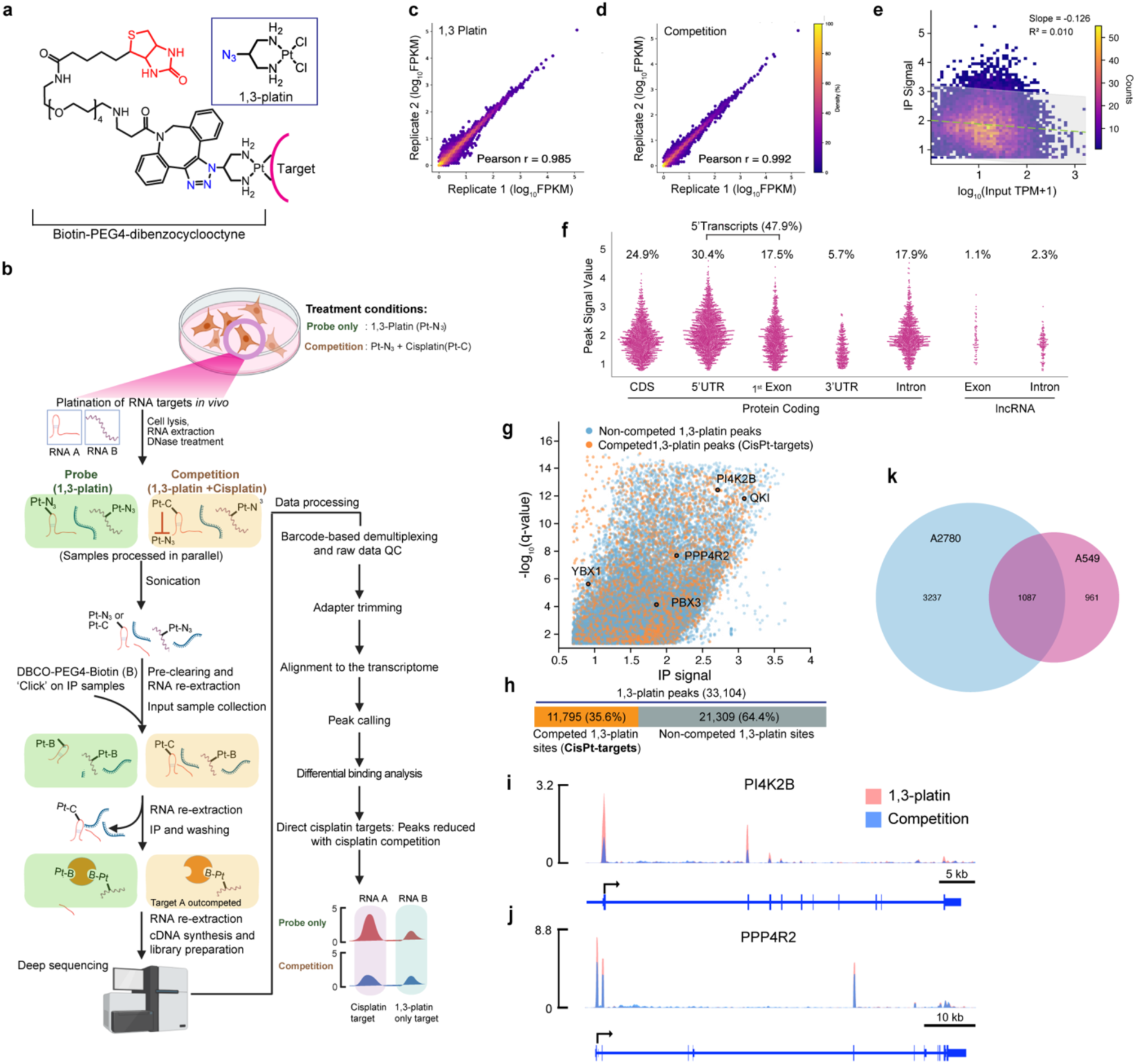
PlatRNA-seq reveals the transcriptome-wide interaction landscape of cisplatin. (**a)** Structure of 1,3-platin and target-bound 1,3-platin conjugated to biotin-PEG4 moiety using click chemistry. (**b**) Workflow schematic of PlatRNA-seq. (**c,d)** Replicate scatter plot of PlatRNA-seq signal (1,3-platin alone) (**c**) or competition samples (**d**) from A2780 cells. The R-value was derived from the Pearson correlation analysis. (**e**) Scatter plot showing PlatRNA-seq pull-down signals versus RNA expression levels in A2780 cells, revealing no correlation between transcript abundance and pull-down efficiency (R2 = 0.01). (**f**) The relative distribution of cisplatin targets across different transcript locations in A2780 cells. (**g)** PlatRNA-seq data visualized as dot plots. Each dot represents a 1,3-platin-interacting RNA fragment identified through peak calling. RNA fragments showing reduced binding (Log₂FC< -0.5) are designated as cisplatin targets (CisPt-targets) and highlighted in orange. Five CisPt-targets, for which read density plots are shown, are circled. (**h)** Quantitation of 1,3-platin–RNA peaks showing the number of sites that were either competed or not competed by cisplatin. (**i, j**) Read density tracks along the transcripts for the genes PI4K2B (**i**) and PPP4R2 (**j**) in A2780 cells. (**k**) Venn diagram of cisplatin targets identified in two cancer cell lines (A2780, A549).

We then measured the IC50 of 1,3-platin in different cancer cell lines to determine treatment concentrations for target identification (Supplementary Fig. 2d). Building on these results, we developed a chemico-transcriptomic method to comprehensively identify platinum-interacting RNAs in mammalian cells (Fig. 2b). The method involves treating the cells with 1,3-platin at IC50 concentration to facilitate binding to cellular platinum-targets. The total RNA is extracted and fragmented by sonication. The purified RNA fragments are subjected to copper-free, bioorthogonal click reaction with DBCO-PEG4-Biotin. The purified RNAs are enriched by streptavidin pulldown, with rigorous washing to reduce background before elution. To specifically map the platinum binding on non-abundant RNAs, rRNA is depleted (optional) during the sequencing library preparation step. The prepared libraries are subjected to high-depth sequencing (See Methods for more details). Reads uniquely mapped to the genome were retained for downstream processing. Deep sequencing data showed strong concordance between replicate experiments (Fig. 2c–d). RNA isolated from pre-enrichment samples was used as a paired input control to identify 1,3-platin-binding peaks using the CLAM peak-calling algorithm ^42^. Parallel competition experiments validated target specificity. Pre-treating cells with cisplatin competitively blocked binding sites, allowing us to identify high-confidence cisplatin-interacting RNA sequences. In this setup, cisplatin occupies its binding sites, blocking access by the clickable analogue 1,3-platin and thereby reducing its binding peak signal, while the signal from binding sites that interact only with 1,3-platin is minimally impacted (Fig. 2b). RNA sites showing at least a 1.5-fold reduction in 1,3-platin enrichment upon competition were classified as cisplatin-interacting regions or cisplatin targets (CisPt-targets) (Fig. 2b).

### Mapping transcriptome-wide interaction of cisplatin

Cellular RNAs may act as a small-molecule sink by spuriously interacting with molecules in a non-specific manner ^43,44^. To address this concern, we investigated whether pull-down efficiency correlated with RNA expression. For RNAs with detectable expression (TPM> 0.1) and IP signal, we found no positive correlation between pulldown efficiency and absolute RNA expression, thereby highlighting the specificity of RNA binding. (Fig. 2e and Supplementary Fig. 3c).

We observed that the 1,3-platin signal was found to preferentially accumulate at the 5’ end of these transcripts, coinciding with the loss of signal upon cisplatin treatment. By overlapping called peaks with key transcript features, we found that ∼48% of cisplatin-target sites overlapped with the 5’ region of transcripts (5’ UTR + first exon) (Fig. 2f). Out of 33104 1,3-platin peaks, 11795 corresponding to 4324 annotated RNAs were competed out upon cisplatin treatment (CisPt-targets) (Fig. 2 g–h). Of these targets, 3,438 have a TPM greater than 1, corresponding to 6.8% of all expressed transcripts (TPM > 1) in A2780 cells. Read density plots of five targets (PI4K2B and PPP4R2, PBX3, QKI, and YBX1) are provided as examples of decreased 1,3-platin binding upon cisplatin treatment (Fig. 2i–j, and Supplementary 2e–g). We also performed PlatRNA-seq in the lung carcinoma cell line A549. A substantial fraction of cisplatin-interacting transcripts was shared between the cell lines, further suggesting specificity of RNA-targets (Fig. 2k). Moreover, the platin–RNA distribution (Supplementary Fig. 3d–j) and other experimental quality metrics (Supplementary Fig. 3a–c) were generally consistent across both cell lines, underscoring the reproducibility and robustness of the assay.

### Cisplatin binds to RNA G-quadruplexes and modifies their biophysical properties

RNA structure plays a crucial role in all stages of the RNA life cycle, including transcription and stability ^45^, and is emerging as a promising therapeutic target ^46,47^. To determine whether platinum drugs preferentially bind specific RNA elements, we performed motif enrichment analysis of platinum–bound peaks from A2780 cells. The most significantly enriched motifs were guanine-rich, suggesting a bias toward G-rich RNA structures. (Fig. 3a). Similar motifs were also enriched in the PlatRNA-seq data from A549 cells (Supplementary Fig. 3k). This result is consistent with the known binding of cisplatin with guanine residues in DNA ^48^ and further validates the specificity of PlatRNA-seq. A prominent RNA secondary structure formed by guanine-rich regions is the RNA G-quadruplex (rG4), a four-stranded RNA structure stabilized by stacked guanine quartets ^49–51^. Using the QGRS Mapper tool ^52^, the top motif identified in A2780 cells and the fourth-ranked motif in A549 cells were predicted to form rG4s (Fig. 3a and Supplementary Fig. 3k, highlighted with red underline).

**Fig. 3.**
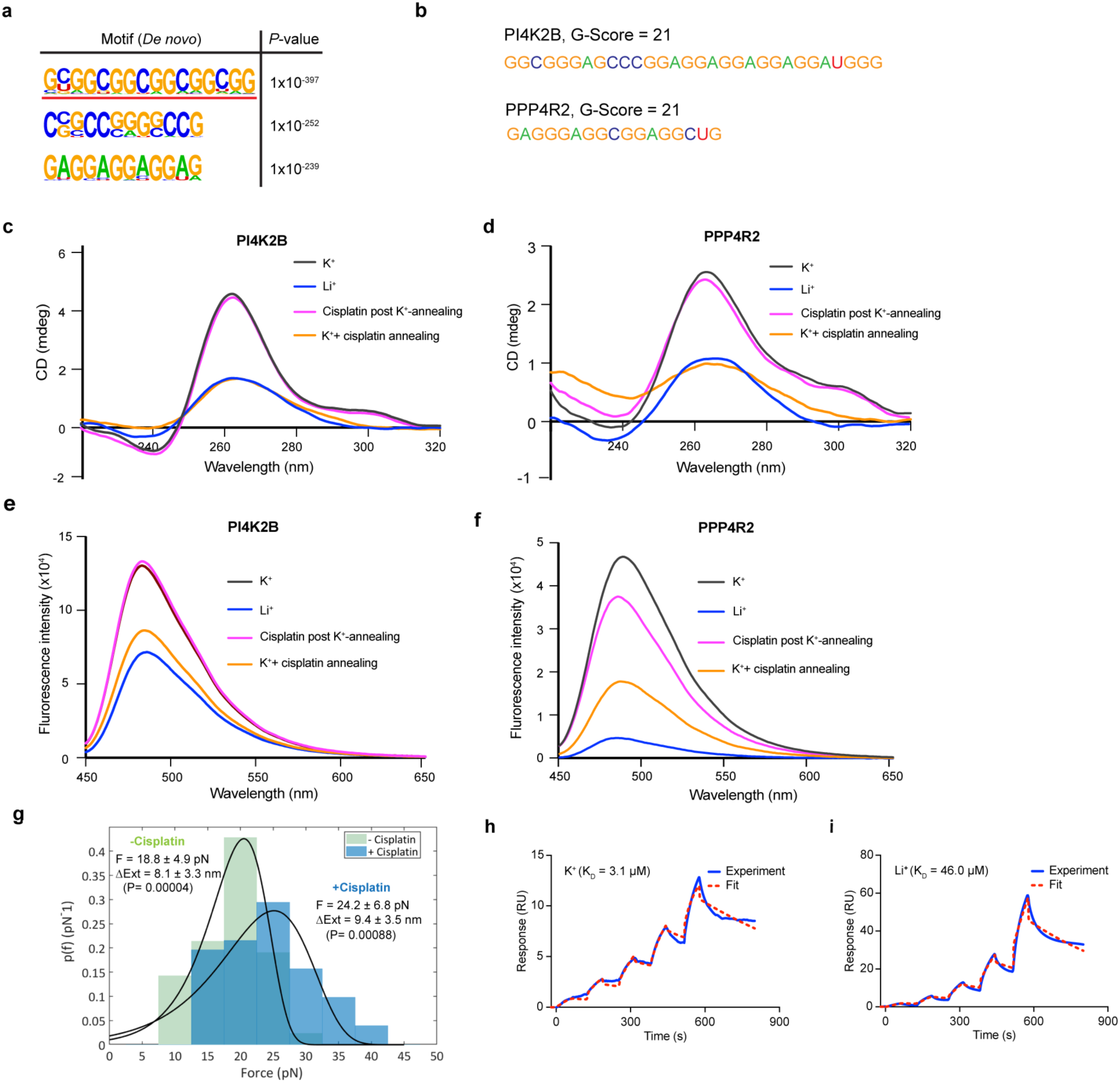
Structural and biophysical characterization of platinum-bound RNAs. (**a)** Motif analysis of cisplatin-target enriched sequences reveals G-rich motifs considering motif lengths of 12, 16, and 20 nucleotides. The top motif underlined in red is a predicted rG4. (**b)** Predicted rG4 sequences from cisplatin-bound regions of the indicated RNAs. G-quadruplex propensity was predicted using QUADRatlas ^59^. A G-score above 19 is considered high rG4 forming propensity. (**c,d)** Circular dichroism spectrum of PI4K2B (**c**) and PPP4R2 (**d**) RNA oligos in rG4-conducive (K+, black) or rG4-disruptive (Li+, blue) buffer conditions, demonstrating their rG4-forming potential. The experiments were also performed under conditions in which cisplatin was incubated with preformed rG4 (purple) or in which annealing was performed in a buffer containing both K+ and cisplatin (orange). (**e,f**) ThT-fluorescence spectroscopy of PI4K2B (**e**) and PPP4R2 (**f**) RNA oligos in rG4-conducive (K+, black) or rG4-disruptive (Li+, blue) conditions. The experiments were also performed where cisplatin was incubated with preformed rG4 (purple) or in which annealing was performed in a buffer containing both K+ and cisplatin (orange). (**g**) Probability distribution of the forces required to unfold PI4K2B rG4 in the absence and presence of cisplatin, measured by optical tweezers (N = 42 for “-cisplatin” and N = 51 for “+ cisplatin”). P-value was calculated using the Kolmogorov-Smirnov test. (**h,i**) SPR dissociation profiles demonstrate cisplatin binds less efficiently to PI4K2B RNA oligo under rG4-disrupting (Li+, KD = 46.0µM) conditions (**i**) compared to the rG4-stabilizing (K+, KD = 3.1µM) buffer condition (**h**).

To experimentally test whether the computationally predicted rG4s contribute to cisplatin-RNA interaction, we selected cisplatin-bound sequences from two RNAs that are identified as cisplatin target from two cell lines (Fig. 2g and Supplementary Fig. 3e), PI4K2B, a PI4 Kinase implicated in invasive cancer phenotype ^53,54^, and PPP4R2, a phosphatase involved in DNA damage repair, both predicted to form rG4 structures by QGRS Mapper ^52^ (Fig. 3b). To experimentally validate rG4 formation, we leveraged the distinct effects of monovalent cations on G-quadruplex stability: potassium ions (K⁺), which promote rG4 folding by stabilizing G-quartet stacking, and lithium ions (Li⁺), which disfavor rG4 structure ^55^. RNA oligonucleotides annealed in K⁺- or Li+-containing buffers were analyzed using circular dichroism (CD) ^56^ and Thioflavin T (ThT) fluorescence^57^. In the K⁺ condition, both sequences displayed rG4-characteristic CD peaks (positive peak near 265 nm, and a negative peak around 240 nm ^58^ (Fig.3c, d), and enhanced ThT fluorescence (Fig.3e,f). These effects were diminished in a Li+-containing buffer (Fig. 3c-f), consistent with the rG4 formation by these sequences under K+ conditions. To assess how cisplatin affects these rG4 structures, we first folded RNA oligos into rG4s in K⁺ buffer before incubating them with cisplatin. CD and ThT data indicated that pre-formed rG4s retained their conformation despite cisplatin treatment. In contrast, when folding was attempted in the presence of both K⁺ and cisplatin, rG4s failed to form, with CD and ThT profiles resembling those in Li⁺ buffer (Fig.3c-f). We next used single-molecule optical tweezers to measure the dynamic mechanical changes in rG4 upon cisplatin binding. Pre-formed PI4K2B rG4 oligo, with or without cisplatin treatment, was unfolded using optical tweezers. The unfolding experiments showed that cisplatin association significantly increased the required force to unfold the rG4 structure by an average of 5.4 pN, from 18.8 ± 4.9 pN to 24.2 ± 6.8 pN (two-sample t-test assuming equal variance and two-sample Kolmogorov-Smirnov test: p = 0.000043 and p = 0.000880, respectively) (Fig. 3g and Supplementary Fig. 4a). This suggests that cisplatin interacts with and stabilizes pre-formed rG4 structures. No significant difference was observed between the change in length upon unfolding (ΔX) in the absence or presence of cisplatin, indicating that cisplatin binding does not change the pre-formed rG4 conformation significantly (Fig. 3g). To evaluate how rG4s influence cisplatin RNA binding affinity, we annealed the PI4K2B rG4 sequence under rG4-promoting (K⁺ buffer) and rG4-inhibiting (Li⁺ buffer) conditions. We then assessed whether the rG4 conformation enhances cisplatin binding with RNA using SPR. Cisplatin exhibited consistently stronger binding to RNA oligos annealed under rG4-promoting conditions, suggesting a higher affinity for rG4-structured RNA (Fig. 3h–i, and Supplementary Fig. 4b). Collectively, these results demonstrate that cisplatin preferentially interacts with and stabilizes existing rG4s, while inhibiting the formation of new rG4s.

### Cisplatin-mediated Pol II stalling and R-loops facilitate 5’-platination of RNA

Guanine-rich RNAs and rG4 can promote the formation of three-stranded DNA-RNA hybrid structures known as R-loops ^60,61^. Given the high affinity of cisplatin to rG4 structures (Fig. 3h), we wanted to examine if the preferential 5’ enrichment of platins is linked to R-loops. We first performed RNA-immunoprecipitation with BG4, a G4-specific antibody (BG4-RIP-seq) ^62,63^, to identify all rG4-harboring RNAs in cisplatin-treated A2780 cells. Comparison with PlatRNA-seq data revealed that cisplatin-target RNAs show higher BG4-RIP enrichment consistent with higher rG4 content (Fig. 4a), consistent with the rG4 enrichment observed in cisplatin-bound RNAs (Fig. 3a). The R-loop CUT&Tag assay ^64^ revealed that the cisplatin-target RNAs exhibit elevated R-loop levels near promoter proximal regions (Fig. 4b–c). Interestingly, the R-loop formation was further increased by treatment with cisplatin (Fig. 4b). These results indicate that cisplatin-target RNAs, enriched in rG4s, preferentially form R-loops near transcription start sites.

**Fig. 4.**
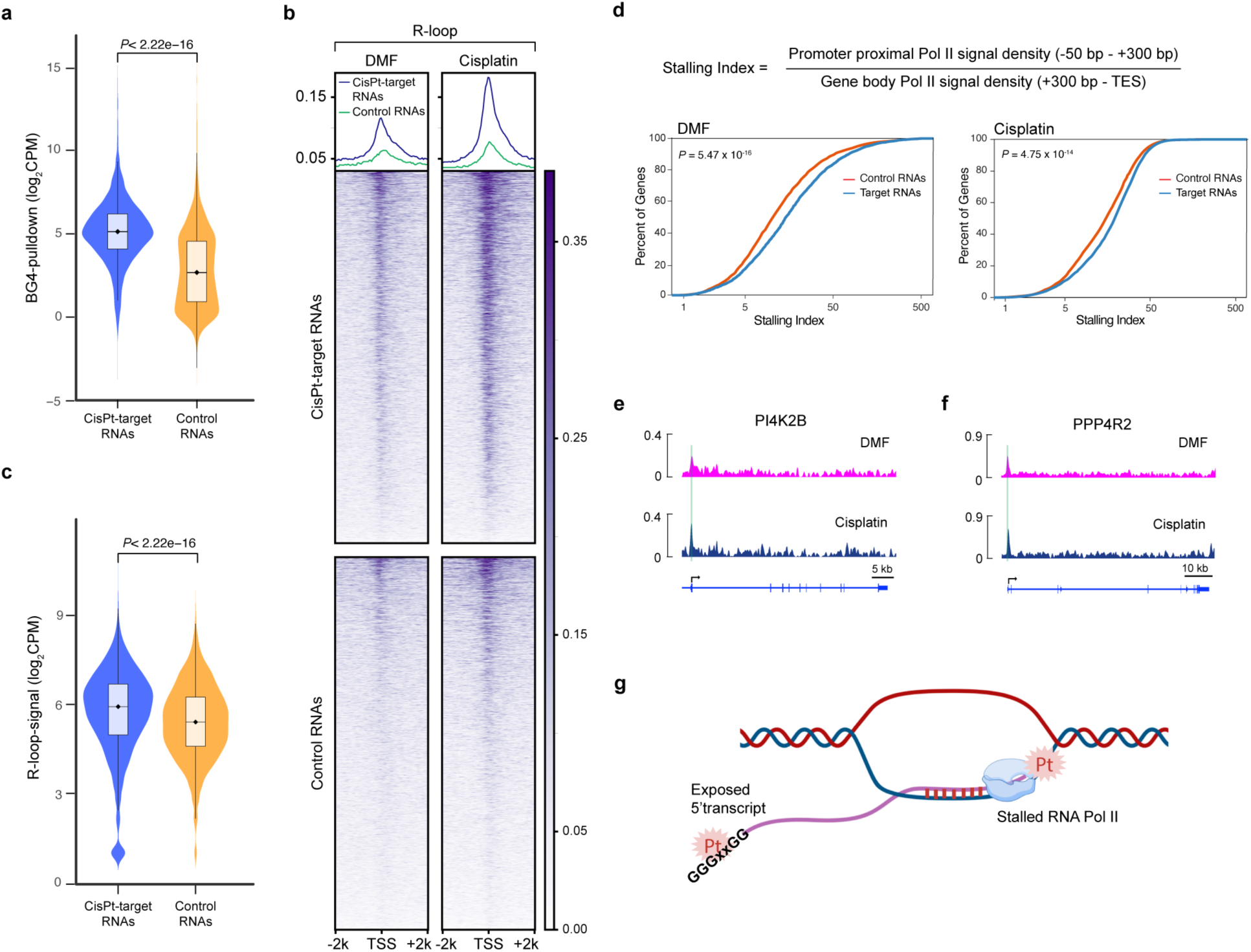
Cisplatin–RNA adducts induce elevated R-loops and transcriptional stalling. (**a**) BG4-RIP-seq data showing higher rG4-signal in cisplatin-target RNAs compared to control RNAs. P-value was calculated using the Wilcoxon Rank-sum test. (**b**) Heatmap showing elevated R-loop in the TSS-proximity of loci coding for cisplatin-target RNAs compared to control RNAs. The R-loop level is further increased upon cisplatin treatment. (**c**) Quantitation of R-loop signal from cisplatin-target and control RNA TSS proximal regions. P-value was calculated using the Wilcoxon Rank-sum test. (**d**) Stalling Index calculation to assess the relative enrichment of RNA Pol II as the gene promoter proximity compared to the rest of the gene body. Cumulative frequency distribution of stalling index reveals elevated promoter proximal stalling of RNA Pol II at cisplatin-target RNAs compared to the control RNAs. P-value was calculated using the two-sample Kolmogorov-Smirnov test. (**e,f**) Read density tracks of RNA Pol II ChIP-seq at PI4K2B (**e**) and PPP4R2 (**f**) genomic loci. The promoter proximal region is highlighted in green. (**g**) Model whereby high rG4 abundance and transcriptional stalling due to cisplatin–DNA adducts lead to elevated R-loops. The exposed 5′ RNA strand enhances the likelihood of cisplatin-RNA interactions.

R-loop accumulation is linked to promoter-proximal pausing of RNA polymerase II ^65^, while DNA crosslinking lesions can induce RNA polymerase II stalling, resulting in proteasome-mediated degradation of the RNA polymerase II complex ^66,67^. Consistent with other reports ^68^, western blot analysis of A2780 cell lysates revealed a marked reduction of Pol II within 3–6 hours post-cisplatin treatment (Supplementary Fig. 5a–b). Therefore, we calculated the stalling index from RNA Pol II chromatin enrichment in response to a 1-hour cisplatin treatment (Fig. 4d). Cisplatin-target genes displayed higher stalling indices under both basal and treated conditions relative to non-targets (Fig. 4d), as demonstrated for several cisplatin targets (Fig. 4e–f and Supplementary Fig. 5c–d). Together, these findings suggest that cisplatin-induced Pol II stalling, coupled with high rG4 content, promotes R-loop formation, exposing 5′ transcript ends for cisplatin binding and leading to preferential platinum enrichment at the 5′ regions of transcripts (Fig. 4g).

### RNA G-quadruplex binding underlies a noncanonical cytotoxic mechanism of cisplatin

Despite the high probability of anticancer small-molecule RNA off-targeting ^4^ (Fig. 1a), the biological and pharmacological roles of RNA off-targeting remain unclear. We sought to determine whether the interaction of cisplatin with rG4 may also contribute to its cytotoxic effects in addition to the DNA damage-induced cell death ^21^. We used two G4 ligands: carboxy-pyridostatin (cPDS), a derivative of the commonly used G4 ligand pyridostatin (PDS) with a binding preference for RNA G-quadruplexes ^69,70^, and PhenDC3 ^71^. Given the high affinity of cisplatin for preformed rG4s (Fig. 3h), we hypothesized that rG4-stabilizing ligands would enhance cisplatin recruitment and cytotoxicity. To test this hypothesis, we pretreated A2780 cells with cPDS followed by cisplatin. Unexpectedly, cPDS pretreatment increased the IC50 of cisplatin (reduced cytotoxicity) in A2780 cells (Fig. 5a), with an average increase of 31.1% (Fig. 5b) in a dose-dependent manner (Supplementary Fig. 6a). The ability of cPDS to inhibit cisplatin activity suggests a competitive interaction between these drugs for rG4 occupancy. Supporting this model, PDS has been reported to stabilize the guanine tetrad plane through π–π stacking and hydrogen bonding with guanine bases ^72^, the same region recognized by cisplatin ^15^. To test this competitive binding hypothesis, we first performed a SPR binding assay using PI4K2B RNA and DNA oligonucleotides. These oligonucleotides were annealed in a K+-containing buffer to induce G4 conformation and immobilized on the SPR chip. The oligonucleotides were then sequentially exposed to cPDS (or vehicle control), followed by cisplatin. Pre-binding with cPDS markedly reduced the cisplatin binding to the RNA oligonucleotides (Fig. 5c), confirming the competitive occupancy of rG4 structures. Consistent with the reported RNA specificity of cPDS ^69,70^, the competitive cisplatin inhibition was RNA-specific, as cPDS pre-treatment did not significantly alter cisplatin binding to DNA G-quadruplex (Fig. 5d).

**Fig. 5.**
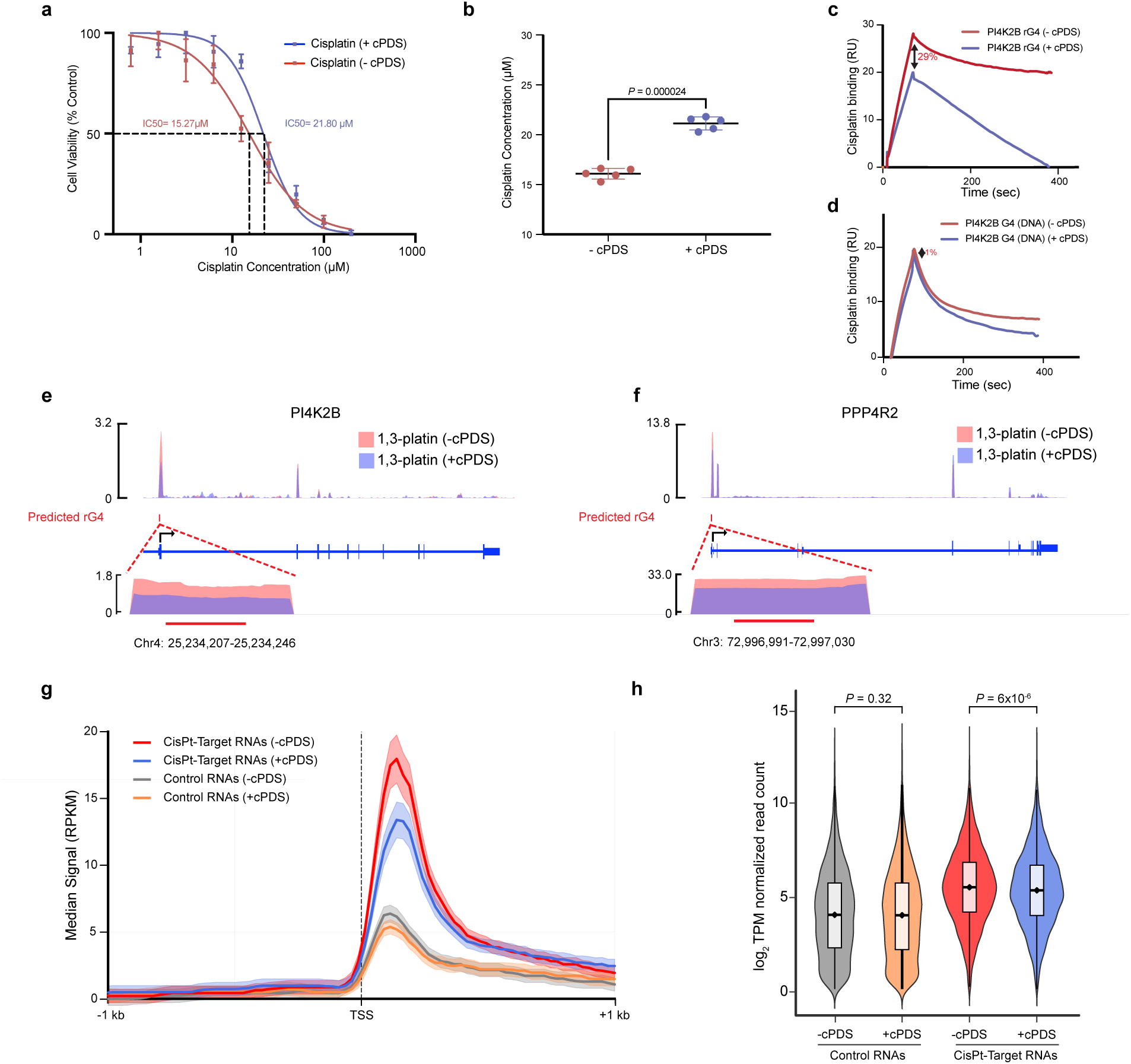
rG4-mediated accumulation of cisplatin promotes cellular cytotoxicity. (**a)** Cell viability assay showing lower efficacy of cisplatin (increased IC50) in cells pretreated with rG4-ligand cPDS. (**b)** Distribution of IC50 change in replicate experiments. P-value was calculated using Paired Student’s t-test. (**c,d**) SPR data showing reduced cisplatin binding to PI4K2B RNA-G4 (**c**) or DNA-G4 (**d**) in the absence (red) or presence (blue) of cPDS pre-treatment. (**e,f**) Read density tracks along the transcripts for the genes, PI4K2B (**e**), and PPP4R2 (**f**) in A2780 cells from 1,3-platin-pulldown sequencing experiments. The cells were treated with vehicle (-cPDS, pink) or pretreated with cPDS (+cPDS, purple) before treatment with 1,3-platin. 1,3-platin signal at the predicted rG4 sites is zoomed in to show decreased 1,3-platin upon cPDS-pretreatment. (**g**), Transcriptome-wide 1,3-platin signal enrichment at the promoter proximal regions of cisplatin-target RNAs (CisPt-Target RNAs) or non-cisplatin target control RNAs in cells pretreated with cPDS (+cPDS) or vehicle (-cPDS). (**h**) Normalized read distribution and quantitation of 1,3-platin signal in the promoter proximal regions of control and cisplatin target RNAs.

To assess this competition in vivo, A2780 cells were pretreated with cPDS or vehicle for six hours before Plat-RNA-seq. cPDS pretreatment markedly reduced 1,3-platin binding at predicted 5′ rG4 motifs in cisplatin-target RNAs (Fig. 5e–f and Supplementary Fig. 6 b–d). Quantification of 1,3-platin peaks at transcript 5′ ends confirmed a significant reduction in cisplatin recruitment to target RNAs (CisPt-target RNAs), but not controls (Fig. 5g). Transcriptome-wide analysis showed a global cPDS-dependent decrease in 1,3-platin binding of CisPt-target RNAs (Fig. 5h). The in vitro and in vivo cPDS competition experiment suggests that this small molecule sterically blocks cisplatin’s access to guanine N7 sites in rG4, not DNA G4, thereby reducing the in vivo association of platinum drugs with structured RNA regions. Consistently, treatment with PhenDC3 also yielded comparable results, diminishing cisplatin-induced cytotoxicity in a dose-dependent manner (Supplementary Fig. 6e–f). Together, these findings suggest that the RNA-dependent cytotoxicity of cisplatin largely arises from its engagement with noncanonical rG4 structures.

## Discussion

Although several FDA-approved small molecules have been predicted to bind RNA ^4^, the prevalence and functional impact of these interactions in clinically used anticancer drugs remain unknown. Our results reveal that a meaningful fraction of compounds present in the library interact with RNA. We used cisplatin, a clinically established chemotherapeutic, as a model to systematically investigate the biochemical basis and functional consequences of RNA off-target interactions. Here, we demonstrate that, in addition to its well-characterized DNA interaction, this drug binds extensively to RNAs, with a strong preference for guanine-rich transcripts capable of forming rG4. We developed PlatRNA-seq, a novel open-ended approach to map cisplatin-interacting RNAs in living cells. We employed this approach and integrated transcriptomic and biophysical analyses to uncover an unexpected, rG4-based mechanism underlying cisplatin-induced cell death, thereby revealing a noncanonical RNA-mediated component of cisplatin action. Beyond cisplatin, these results illustrate the functional impact of unintended RNA binding by clinically used small-molecule drugs, and our strategy provides a roadmap for the systematic characterization of the biochemical basis and functional consequences of RNA off-targeting.

To our knowledge, PlatRNA-seq is the first open-ended approach for transcriptome-wide mapping of RNA targets of a clinically used small molecule. While analogous methods exist for cisplatin-DNA adducts ^32,34,73^, no comparable strategy has been applied to RNA. Several lines of evidence argue that cisplatin-RNA binding is non-random. First, cisplatin binds only to a small fraction (6.8%) of the total cellular RNAs. Second, cisplatin-RNA binding is independent of transcript abundance (Fig. 2e and Supplementary Fig. 3c). Third, interactions are enriched at the 5’ transcript ends. Fourth, cisplatin-bound RNA sequences are preferentially enriched for rG4 motifs. We focused on the cisplatin-rG4 interaction, given their higher binding affinity and the availability of chemical tools for mechanistic investigation^49^.

Our study is the first to characterize cisplatin-rG4 binding and to reveal a functional role in cytotoxic mechanisms. Our biophysical data reveal a dichotomous effect of cisplatin on rG4 conformation. When the rG4 conformation was induced by annealing the oligos in the presence of both K+ and cisplatin, the canonical rG4 pattern was not observed in both CD and fluorescence spectroscopy, indicating that stable rG4 failed to assemble. In contrast, optical tweezer experiments revealed a stabilizing effect of cisplatin on preformed rG4s. We propose that in the unfolded state, guanine N7 platination and crosslinking sterically disrupt G-tetrad stacking and prevent rG4 formation. In preformed rG4s, spatial proximity of juxtaposed guanines within the folded topology increases their availability for crosslinking, thereby enhancing the affinity of cisplatin for rG4s and their thermodynamic stability. In support of this model, prior data demonstrate that cisplatin modulates the folding kinetics of DNA G-quadruplex^74^. Platinum-DNA-G4 interactions have shown promising value in therapeutic applications through cisplatin-pyridostatin conjugates ^75^ and cisplatin-terpyridine complexes, which direct platinum drugs to DNA G4 and improve therapeutic efficacy and reversal of chemoresistance. Based on the contribution of RNA-cisplatin binding in cytotoxicity (Fig. 5a, b), conjugating platinum to carboxy-pyridostatin^69^ or other rG4 selective ligands offers a promising alternative strategy to improve cisplatin efficacy and address the persistent challenge of platinum chemoresistance^11^.

G-quadruplexes (G4s) are increasingly recognized as therapeutic targets in oncology ^76–78^. Several G4 ligands induce DNA damage responses and enhance the efficacy of chemotherapeutic agents in preclinical models ^79–82^. These observations have contributed to an expectation that G4 ligands act broadly synergistically with standard chemotherapies. However, our data indicate that G4 ligands and chemotherapy agents can also exhibit antagonistic interactions in a concentration and treatment regimen-dependent manner. Several factors likely underlie these divergent outcomes. G4 structures are cell-type specific and shape lineage-specific transcriptional programs ^83–85^. For example, sublethal PDS resensitizes resistant, but not parental, cells to cisplatin by modulating a resistance-promoting transcriptional program ^86^. Synergy between chemotherapeutics and G4-ligands also depends on treatment regimen, dosage, and ligand choice. A recent study combining cisplatin with rG4 ligands such as PDS, Berberine, and RHPS4 observed synergistic effects in a minority of experimental settings, whereas most combinations were antagonistic ^87^. Notably, when cells were pretreated with G4 ligands, a treatment regimen employed in this study, all three antagonized cisplatin. Berberine and PDS yielded combination indices (CI) of ∼1.28 and 1, respectively, and RHPS4 reduced cisplatin efficacy by more than 50%. Other drugs with potential RNA-related mechanisms, including doxorubicin ^88,89^ and 5-fluorouracil ^90^, exhibit similar G4-ligand-mediated antagonism ^87^. Together, these findings suggest that while rG4 interactions with chemotherapeutic agents are a functional mechanism of action for certain anticancer drugs, antagonism may also be an outcome if dose and timings are not carefully calibrated, particularly among drugs with rG4-binding-related functions. Appreciating this complexity will be key to advancing rG4-targeted therapies and rational drug combinations.

However, it is possible that cisplatin binding to non-rG4 sequences or adenines may also exert biological effects by generating RNA adducts, leading to translational arrest or activation of RNA damage response pathways. Cisplatin and other DNA-binding chemotherapeutics have been shown to induce translation defects and ribosome stalling ^91,92^. In addition, emerging evidence suggests that RNA damage response (RDR) pathways are important mediators of the cellular stress induced by environmental and chemotherapy agents ^93,94^. Distinct RNA lesions engage specific stress responses. For instance, RNAs damaged by UV-radiation are processed within stress granules enriched for the RNA helicase DHX9 ^95^, while 5-fluorouracil–incorporated RNAs trigger RDR during ribosome biogenesis ^90^. rG4 structures exert multifaceted functions throughout the RNA life cycle ^49,51^, including stability, translation, and decay, by recruiting RNA-binding proteins (RBPs) ^50^. A systematic investigation into how oncology small molecules identified in our RNA-binding screen (Supplementary Fig. 1c) modulate RBP–rG4 interactions and thereby alter the stability and fate of target RNAs will uncover novel mechanistic insights into the RNA-dependent pharmacology of cisplatin and other anticancer drugs.

RNA off-targeting among anticancer small molecules is likely far more prevalent than our findings suggest (Supplementary Fig 1c-d) due to several inherent limitations of our experimental approaches and assay conditions. First, we employed only two simplified RNA pools (a random 20-mer with unknown composition, and an equimolar mixture of homopolymer RNA). This will likely undercount the structure-dependent RNA-small-molecule interactions that may arise under the diverse physiological conditions of the intracellular environment. Second, binding reactions were conducted under a single defined set of assay conditions, precluding the detection of interactions that depend on specific RNA secondary structures, ionic environments, and other molecular factors present in vivo. Third, our screen covered only ∼40% of the FDA-approved anticancer drugs cataloged in DrugBank (please see methods). Future studies employing an exhaustive panel of clinically used anticancer agents screened against an array of RNA species representative of the structural and compositional complexity of the transcriptome will be essential for generating a comprehensive picture of RNA off-targeting. Despite these limitations, our study offers several important contributions. Our screening strategy represents an accessible and reproducible platform for conducting RNA off-target screening of small molecules. The observation that a meaningful fraction of drugs in our limited library bind to RNA provides the first experimental glimpse into the magnitude of RNA off-targeting among clinically approved anticancer agents.

In conclusion, our study reveals widespread RNA binding by anticancer drugs, including platinum agents, redefining RNA as a biologically relevant chemotherapeutic target. These findings highlight the potential to design next-generation platinum compounds with enhanced selectivity for rG4s, thereby overcoming current limitations of this drug class ^96,97^. More broadly, the prevalence of RNA interactions among widely used anticancer agents reveals a hidden layer of drug activity, underscoring the need to integrate RNA into drug design and resistance studies.

## Supplementary Data

Supplementary Table is provided separately

## Data Availability

The GEO accession ID for processed and raw next-generation sequencing data is GSE310425.

## Acknowledgments

The authors acknowledge the support of Dr. Steven Metallo, Department of Chemistry, for assistance with the circular dichroism experiments, Dr. Bogdan Tanasa (Sofia Informatics), and Dr. Robert Suter, Lombardi Comprehensive Cancer Center, for consultation on data analysis. We thank the National Cancer Institute Developmental Therapeutics Program (NCI/DTP) for providing the small molecule library (AOD XI) used in this study. The Biacore Molecular Interaction Shared Resource (BMISR) facility at Georgetown University. The BMISR is supported by NIH grant P30CA51008 and 1S10OD019982–01. ICP-MS measurements were performed in the OHSU Elemental Analysis Core with partial support from NIH (S10OD028492). This study was supported by the Department of Defense (OC220133) to SJN and VJD, NSF (CHE2109255) to VJD, and NIH (R35GM150636) to SJN.

## Author Contributions

A.K., W.G., and R.T. performed and analyzed PlatRNA-seq. X.W. conducted biophysical assays. A.H., A.K., and R.M. conducted optical tweezer experiments and analyzed the data. X.W., A.K., S.B.S., P.B.T., and A.U. performed SPR assays and analyzed the data. W.G. and R.T. performed and analyzed BG4-RIP-seq and R-loop CUT&Tag experiments. S.B.S. and R.T. performed and analyzed the ChIP-seq experiment. A.K. performed ICP-MS experiments. X.Z. analyzed the small molecule database. W.G., X.W., and S.B.S. performed cell assays with assistance from A.F. and D.K. K.R.A. conducted chemical synthesis. V.J.D. contributed to chemical formulation, experimental design, and data interpretation. S.J.N. conceived and supervised the project, designed the experiments, wrote the initial manuscript, and secured funding. All authors contributed to data interpretation and manuscript writing.

## Materials and Methods

### Cell culture

Human ovarian adenocarcinoma cell line A2780 was obtained from Sigma-Aldrich. A549 lung adenocarcinoma cell line and TOV112D ovarian cancer cell lines were obtained from Georgetown University’s Tissue Culture and Biobanking Shared Resource. A2780 and TOV112D cells were maintained in RPMI supplemented with 10% FBS and 1% penicillin–streptomycin at 37 °C with 5% CO₂. A549 cells were maintained in DMEM supplemented with 10% FBS and 1% penicillin–streptomycin under identical conditions. All cell lines were passaged every 3–4 days and periodically tested to ensure the absence of mycoplasma contamination.

### PlatRNA-seq assay

A2780 cells were treated in two biological replicates with the IC₅₀ concentrations of either 1,3-platin or dimethylformamide (DMF) for 16 hours. For competition experiments, cells were first incubated with 10 μM cisplatin for 8 hours and subsequently treated with 1,3-platin for an additional 16 hours. Total RNA was extracted using TRIzol (Thermo Fisher, MAN0001271) according to the manufacturer’s instructions and resuspended in 100 μL of nuclease-free water, and subjected to sonication (Diagenode Bioruptor, high power) for three cycles of 30 seconds on/30 seconds off at 4 °C. Streptavidin Dynabeads T1 (Thermo Fisher, 65601) were prepared by washing three times with Bead Binding & Wash Buffer I (1 M NaCl, 10 mM Tris-HCl pH 7.5, 1 mM EDTA), twice with Solution A (0.1 M NaOH, 0.05 M NaCl), and twice with Solution B (0.1 M NaCl), followed by resuspension in Buffer I (2 M NaCl, 20 mM Tris-HCl pH 7.5, 2 mM EDTA). To remove nonspecific bead binders, equal volumes of washed beads and sonicated RNA (100 μL each) were incubated together for 30 minutes at room temperature with gentle rotation. The supernatant fraction was collected, and RNA was re-extracted using TRIzol LS (Thermo Fisher, MAN0000806) and resuspended in 100 μL. Ten percent of each lysate was reserved and stored at -80 °C as an input control.

90 μL of pre-cleared RNA was combined with 100 µM DBCO-PEG4-Biotin (Vector Laboratories, CCTA1055) and SUPERase•In RNase inhibitor (Invitrogen, AM2694; 1:40 dilution), brought to a final reaction volume of 100 µL, and incubated for 16 hours at 37 °C. Following labeling, samples were treated with 2 µL of TURBO DNase (Invitrogen, AM2239) in 11 µL of 10x TURBO DNase buffer at 37 °C for 1 hour. RNA was purified with TRIzol LS (Thermo Fisher, MAN0000806). Biotin-labeled RNA was incubated with freshly prepared, RNase-free Streptavidin Dynabeads T1 (Thermo Fisher, 65601; prepared as described above and supplemented with 100 µg/ml BSA) for 30 minutes at room temperature with gentle rotation. Bead–RNA complexes were washed 12 times: six washes with 1 ml of pre-warmed (65 °C) Bead Wash Buffer II (5 mM Tris-HCl pH 7.5, 0.5 mM EDTA, 1 M NaCl), followed by six washes at room temperature. After the final wash, RNA was eluted from the beads using TRIzol (Thermo Fisher, MAN0001271). To facilitate pellet visualization during isopropanol precipitation, 2 µl of GlycoBlue (Invitrogen, AM9515) was added prior to precipitation. RNA pellets were resuspended in 20 µl of nuclease-free water and quantified using the Qubit RNA HS Assay Kit (Invitrogen, Q32852). Pulldown and input RNA samples were depleted of rRNA and processed for the construction of a stranded cDNA library using the KAPA RNA HyperPrep Kit with HMR RiboErase (Roche, KR1351 v4.21). Library quality was assessed using the TapeStation D1000 High Sensitivity assay. Libraries were sequenced on an Illumina platform, generating ∼50 million paired-end 150 bp reads per sample.

### PlatRNA-seq with cPDS treatment

A2780 cells were cultured in 10 cm dishes and processed in biological replicates. Cells were treated with 30 μM 1,3-platin or DMF for 16 hours. The cPDS (MedChemExpress, HY-112680A) competition samples were incubated with 5 μM cPDS for 8 hours, followed by treatment with 30 μM 1,3-platin for an additional 16 hours. Subsequent steps for RNA isolation, click-chemistry labeling, library preparation, and sequencing were performed as described above for PlatRNA-seq.

### PlatRNA-seq data analysis

Paired-end FASTQ files were quality-checked using FastQC (v0.11.9). Sequencing adapters were trimmed using CutAdapt (v4.0) in paired-end mode. Trimmed reads were aligned to the hg38 reference genome using STAR (v2.7.11), retaining uniquely aligned reads ^98^. Enriched transcripts relative to the input were identified by applying the peak-calling algorithm CLAM (v1.2.3) to the mapped read ^42^. CLAM preprocessing was performed by retagging aligned reads to the median mapped location, using a window size of 50 nucleotides (nt) and a maximum of one tag per position. RNA peaks were identified and annotated using CLAM with a bin size of 100 nt, requiring a minimum pulldown coverage of 4 reads and a fold change over the input of 2 for A2780 datasets, and 6 for A549 datasets. Differential binding analysis was performed using DiffBind (v3.16.0), with read counting over full peaks (summits = 0), and contrasts defined between 1,3-platin-treated and competition-treated samples (two replicates each) ^99^. Differentially bound sites were called using cutoffs of FDR < 0.01 and log2 fold change > 0.5. Downstream analysis was performed using a custom R script. Identified peaks were uniquely assigned to transcripts based on maximum reciprocal overlap.

### Motif search

De novo motif analysis was performed on peaks lost upon cisplatin competition (cisplatin-bound RNA regions) using HOMER (Version 5.1) ^100^, using the following command: findMotifsGenome.pl with the parameters -size 100 -bg -len 12,16,20 -rna -p 24. De novo motifs were computed against a shuffled background of sequences from a DMF-treated control.

### SPR screening of small molecule library for RNA binders

A library of FDA-approved oncology drugs (AOD XI) was obtained from the National Cancer Institute Developmental Therapeutics Program (NCI/DTP). All assays were performed under stringent RNase-free conditions. Bench surfaces, pipettes, and instruments were decontaminated with RNaseZAP, and instrument tubing was sequentially flushed with RNaseZAP for 7 minutes and then with DEPC-treated ultrapure water for 7 minutes.

The screening experiments were conducted using a Biacore 4000 instrument with a CM5 sensor chip at 25 °C. Flow spot (S)3 of all flow cells (FCs) was used as the reference for S1, S2, S4, and S5. Neutravidin (10 mg/ml stock) was diluted in 10 mM acetate buffer (pH 4.5) and immobilized on all four FCs to a level of ∼15,000 RU using standard amine coupling chemistry. Two distinct biotin-tagged RNA pools were diluted in RNA buffer (20 mM sodium phosphate, pH 7.4, 0.05% v/v Tween-20) and captured onto different sensor chips at two different ligand densities. The random 20-mer pool was captured onto S1 of all FCs to a level of ∼1300 RU, and S2 of all FCs to a level of ∼400 RU. The equimolar mixture of 12-mer RNA homopolymers was prepared as a 1:1:1:1 molar ratio of 12bio-rA-mer, 12bio-rG-mer, 12bio-rC-mer, and 12bio-rU-mer, and captured onto S5 of all FCs to a level of ∼250–380 RU, and on S4 of all FCs to a level of ∼100–250 RU. RNA buffer was used as both the immobilization and capture running buffer.

Small molecules (SMs) were designated as hits if they: (i) exhibited the expected dose-response relationship, and (ii) achieved a relative binding response exceeding 25% of theoretical Rmax for the compound. The theoretical Rmax is the maximum expected binding signal, assuming 100% of the ligands on the surface are occupied by a single analyte. Since the binding signal is proportional to how much ligand is on the surface and the molecular weight ratios of the analyte to ligand, theoretical Rmax values were calculated for each compound and used for binding decisions. For practical purposes, we employed a 1:1 binding model for these calculations. Compounds that showed more than 100% Rmax may be binding to more than one site on the given RNA oligo or may be binding as multimeric molecules to a single site as opposed to a monomeric 1:1 interaction.

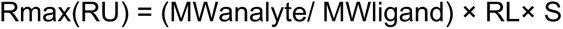

Where, MWanalyte = molecular weight of SM (Da), MWligand = molecular weight of RNA (Da), RL = Immobilized RNA level (Response Units, RU) on the flow cell, S = binding stoichiometry (assuming 1:1). SMs were diluted to final concentrations of 50 µM, 10 µM, and 2 µM in the RNA buffer+5% DMSO. Both contact and dissociation times were set to 60 seconds, and the flow rate was maintained at 30 µL/min. A single 20-second pulse of 1 M NaCl was used for surface regeneration. All analytes were injected once.

Double reference subtraction was applied to account for background signal: (1) The signal from buffer alone (no small molecules) injection on all flow cells was subtracted from the recorded binding values. (2) For each small molecule, the signal from S3 (no RNA surface) on the corresponding FC was also subtracted from the binding signal. If the binding signal did not follow a dose-response pattern among the tested three doses, those compounds were eliminated from further analysis, and the signal was considered non-specific. For the remaining compounds, those that showed a binding signal equal to or greater than 25% and less than 200% of the calculated Rmax were designated as RNA binders.

### SPR assay for cisplatin binding to RNAs

Experiments were performed on a Biacore T200 instrument using CM5 sensor chips at 25 °C. Out of four independent flow cells (FCs), the first FC (FC1) served as the reference for FC2–FC4. Neutravidin was immobilized onto all FCs using the same amine-coupling procedure described in the screening experiment. Biotinylated RNA homopolymers were then captured (∼680 RU) on the chip surface. RNA buffer served as both immobilization and capture running buffers. Cisplatin was injected in two-fold serial dilutions from 40 µM to 2.5 µM in RNA buffer supplemented with 5% (v/v) DMSO. Each concentration of cisplatin was injected in a single-cycle mode with 60 seconds of association and 600 seconds of dissociation at a flow rate of 50 µL/min. Sensorgrams were analyzed via fitting to the 1:1 binding model to calculate kinetic parameters.

### SPR experiments for comparison of cisplatin binding to rG4 and linear RNA

Biotinylated PI4K2B RNA (IDT, 10,392.5 Da) was prepared at 5 µM in either annealing buffer (10 mM Tris, pH 7.5, 100 mM potassium chloride) or annealing-prevention buffer (10 mM Tris, pH 7.5, 100 mM lithium chloride), aliquoted into 0.2 ml PCR tubes, and thermally processed to promote or inhibit G-quadruplex folding. Samples were heated to 95 °C for 5 min in a thermocycler, then cooled to 4 °C at a ramp rate of 0.1 °C/s. SPR experiments were conducted using a Biacore T200 instrument with CM5 sensor chips at 25 °C. FC1 served as the reference for FC2, and FC3 served as the reference for FC4. Neutravidin was immobilized to a level of ∼19,500 RU. PI4K2B folded in annealing buffer was then captured onto FC2 (∼3,650 RU) in RNA buffer-K (10 mM Tris, pH 7.5, 100 mM potassium chloride, 0.05% Tween-20), while PI4K2B folded in annealing-prevention buffer was captured onto FC4 (∼3,800 RU) in RNA buffer-L (10 mM Tris, pH 7.5, 100 mM lithium chloride, 0.05% Tween-20). Cisplatin was then injected at a flow rate of 50 µL/min in single-cycle kinetics mode at concentrations ranging from 40 µM to 2.5 µM (two-fold dilutions) in the appropriate running buffer supplemented with 5% DMSO (RNA buffer-K for FC1/FC2 and RNA buffer-L for FC3/FC4).

### Structural similarity assessment of small-molecule RNA-binding

A comprehensive list of FDA-approved anticancer agents was obtained from DrugBank (v5.5.13, release date January 2, 2025). The list of small-molecule RNA binders was compiled from the R-BIND 2.0 repository ^19^ (accessed January 15, 2025) and the ROBIN database ^20^. Compound CIDs were generated via the PubChem Exchange Service. Substructure key-based, two-dimensional Tanimoto coefficients (Tc) were calculated using the PubChem Score Matrix Service. In total, 1701 CIDs were included (159 from R-BIND 2.0 and 1542 from the ROBIN database), and pairwise Tc values were computed for each. Compounds with Tc values exceeding 0.85 were classified as potential RNA binders.

### Inductively Coupled Plasma Mass Spectrometry (ICP-MS)

The experiment was performed according to a published protocol ^28^. Briefly, adherent cells (3 × 10^5^ per well, 6-well plates) were grown to ∼70–90% confluence and treated with 10 µM cisplatin for 24 hours. Whole-cell samples were collected by trypsinization, washed with PBS, resuspended in HBSS, and counted. Genomic DNA was isolated using the PureLink Genomic DNA Mini Kit (Thermo Fisher, K182001) with RNase A treatment, and RNA was isolated using the Zymo QuickRNA MiniPrep Kit (Zymo Research, R1054) with on-column DNase I digestion, following the manufacturer’s protocols. DNA and RNA were quantified by NanoDrop. DNA and RNA samples were acidified with 70% OmniTrace nitric acid (Sigma-Aldrich, NX0404-1) and digested at 90 °C for 45 min, cooled, and stored at -80 °C. Elemental analysis was performed on an Agilent 8900 ICP-MS (RF 1550 W, plasma 15 L/min, carrier 0.9 L/min, He 5.0 mL/min, KED mode). Platinum and phosphorus were quantified against external standards and normalized to cell counts, nucleic acid mass, or platinum/phosphorus ratios to assess nucleic acid accumulation.

### In vitro Click-Chemistry and Dot blot protocol

Bulk RNA was purified from ∼90% confluent A2780 cells (10-cm plates) using the Zymo QuickRNA MiniPrep Kit (Zymo Research, R1054). Contaminating DNA was removed with on-column DNase I digestion. RNA was eluted in nuclease-free water and quantified by NanoDrop. Aliquots (100 µL) were sonicated (3 × 30s/30s, 4 °C, Diagenode Bioruptor) and incubated with 10 µM 1,3-platin in 100 µL reactions containing SUPERase In RNase inhibitor at 37 °C for 8 hours, protected from the light. The modified RNA was re-purified using the Zymo RNA Clean & Concentrator-5 kit (Zymo Research, R1013) and quantified using a NanoDrop. For click-labeling, 49 µL of platinated RNA was incubated with 1 µL SUPERase RNase inhibitor and 5.5 µL AZDye488-DBCO (Vector Laboratories, CCT-1278; 1 mM in DMSO) at 37 °C for 16 hours in the dark. Labeled RNA was re-purified, eluted in nuclease-free water, and quantified by UV-Vis spectroscopy. RNA (1 µg/µL) was heat-denatured (95 °C, 3 min), spotted onto Amersham Hybond-N+ nylon membranes, UV-crosslinked (254 nm, 125 mJ/cm2), washed (PBS + 0.1% Tween-20), and imaged for AZDye488 fluorescence. Membranes were subsequently counterstained with methylene blue, destained, and re-imaged.

### Optical Tweezers Protocol and Analysis

The rG4 forming sequence from PI4K2B (Fig. 3b, underlined below) was modified with the DNA annealing handles for the optical tweezer experiment (rCrArUrUrGrCrGrCrArUrUrGrCrGrUrUrUrUrGrGrCrGrGrGrArGrCrCrCrGrGrArGrGrArGrGrArGr GrArGrGrArUrGrGrG rUrUrUrGrGrUrArUrArGrCrArUrUrCrUrArG).

Nucleotide handles were annealed to the RNA oligonucleotides (10 µM) using hairpin labeling and tethering kit (Lumicks, SKU: 00016) following the manufacturer’s instructions. 10 µM oligo stock was aliquoted into PCR strip tubes (25 µL per tube) and diluted on ice with an equal volume of either 10 mM Tris (pH 7.5) containing 200 mM KCl or 10 mM Tris (pH 7.5). Samples were annealed by heating to 100 °C for 5 min, followed by cooling to 40 °C at a rate of 1 °C/s. For platination, annealed oligos were incubated with 50 µL of 5 µM cisplatin in 10 mM Tris (pH 7.5) for 16 hours at 37 °C.

Single-molecule force spectroscopy was performed using a MiniTweezers instrument ^101^, following modified force-ramp protocols for rG4 unfolding with or without cisplatin ^102,103^. The microfluidic chamber was equilibrated with rG4 buffer (10 mM Tris, 100 mM KCl, pH 7.5). rG4 samples were incubated with anti-digoxigenin (Lumicks, SKU: 00016) beads for 30 min at room temperature and diluted in 1 mL of rG4 buffer. Platinated rG4 samples (+cisplatin) and untreated rG4 samples (−cisplatin) were processed identically thereafter. Streptavidin-coated beads (Lumicks, SKU: 00016) were diluted in 1 mL of rG4 buffer and injected into the chamber. For tether formation, a streptavidin-coated bead was trapped on the micropipette, and an anti-digoxigenin bead carrying the rG4 was captured in the optical trap. rG4s were unfolded at a pulling velocity of 50 nm/s with a 30 s refolding interval. A total of 51 unfolding trajectories (+cisplatin) and 42 trajectories (−cisplatin) were recorded. Data were analyzed with a custom MATLAB script to extract unfolding forces ^104^. Statistical differences between +cisplatin and −cisplatin conditions were assessed using two-sample t-tests assuming equal variance (p = 0.000043) and two-sample Kolmogorov-Smirnov (KS) (p = 0.000880).

### BG4-RIP-seq

A2780 cells were treated with 25 µM cisplatin for 6 hours in replicates. The pulldown experiment was performed using the RIP kit (MBL Life Science, RN1005) following the manufacturer’s instructions with several modifications, as described below. After treatment, each replicate was processed independently for BG4 pulldown. For each sample, 25 µL of 50% M2 bead slurry (Sigma-Aldrich, M8823) was washed three times with nuclease-free PBS, rinsed once with 500 µL of the wash buffer included in the RIP kit, and then resuspended in 500 µL of lysis buffer (RIP kit) on ice. The cells were collected, washed twice in ice-cold PBS, and pelleted at 300 × g for 5 min at 4 °C. The cell pellets were resuspended in 500 µL of lysis buffer, vortexed, and then incubated for 10 min at 4 °C. Afterward, the samples were centrifuged at 12,000 × g for 5 min. 500 µL of cell lysate was incubated with M2 beads for 1 hour at 4 °C. The supernatant was then collected and incubated overnight at 4 °C with 15 µg of BG4 antibody (Anti-DNA/RNA BG4, scFv fragment, FLAG-tagged, Absolute antibody, Ab00174-30.126). The following day, 50 µL fresh M2 bead slurry was washed as above. Lysates were incubated with beads for 2 hours at 4 °C. The beads were then washed three times with 1 mL of wash buffer. Bead-bound RNA was extracted using the TRIzol method. The rRNA-depleted stranded libraries were prepared with the KAPA RNA HyperPrep Kit with HMR RiboErase (Roche, KR1351 v4.21). Library quality was assessed using the TapeStation D1000 High Sensitivity assay. The libraries were sequenced at a depth of 40 million reads.

### BG4-RIP-seq analysis

Paired-end samples were aligned to transcripts using the comprehensive gene annotation from GENCODE (Ensembl 112, GRCh38.p14) with STAR ^98^. Gene-level read counts were computed with featureCounts from Subread (v2.1.1) ^105^. Read counts were normalized to transcripts per million (TPM). Only transcripts with abundance ≥ 1 TPM in both replicates were retained for downstream analysis. RNA abundance in cisplatin-treated replicates for target versus control RNAs was visualized by normalizing to log2 CPM reads using edgeR (v4.6) ^106^.

### Genome-wide R-Loop mapping

A2780 human ovarian carcinoma cells were cultured under standard conditions and seeded to approximately 70% confluency prior to treatment. Cells were exposed to 25 µM cisplatin for 6 hours, or an equal volume of vehicle (DMF) in biological replicates. Following treatment, cells were collected by gentle scraping in PBS, washed once, and pelleted for subsequent R-loop profiling. Genome-wide mapping of R-loops was performed using the CUT&Tag-IT R-loop Assay Kit (Active Motif, 53167) according to the manufacturer’s instructions, with minor modifications. Briefly, 2–5 × 10^5^ treated cells were resuspended in wash buffer containing protease inhibitors, washed, and immobilized on Concanavalin A-coated magnetic beads. Bead-bound cells were incubated in Complete Antibody Buffer supplemented with digitonin and IGEPAL. To generate RNase controls, 5 µL of RNase A was added to selected samples, while water was added to parallel control samples. Cells were incubated overnight at 4 °C with the mouse monoclonal S9.6 DNA–RNA hybrid antibody. Cells were washed in wash buffer and incubated with anti-mouse secondary antibody for 2 hours at room temperature. After additional washes, cells were exposed to pre-assembled pA–Tn5 transposomes (1:100 dilution in Dig-300 Buffer) for 1 hour at room temperature, washed, and subjected to tagmentation in Complete Tagmentation Buffer at 37°C for 1 hour. The reaction was terminated with EDTA, SDS, and Proteinase K, and DNA fragments were released by incubation at 55°C for 1 hour. DNA was recovered using silica spin column purification with the reagents provided in the CUT&Tag-IT R-loop Assay Kit and eluted in 22 µL of elution buffer. Sequencing libraries were prepared directly from eluted DNA using the reagents provided in the kit, following the manufacturer’s instructions. Libraries were amplified using Q5 polymerase, following the cycling steps: 72°C for 5 minutes, followed by (98°C for 10 seconds, 63°C for 20 seconds) × 12 cycles, and a final extension at 72°C for 1 minute. PCR products were purified by a double-sided SPRI bead selection (ratio of 0.5x followed by 1.2x) to remove large fragments and size-select for library inserts. Final libraries were eluted in 20 µL of elution buffer, quantified by Qubit fluorometry, and assessed for quality by TapeStation D1000 High Sensitivity assay. Pooled libraries were sequenced on an Illumina platform to generate paired-end 150-bp reads, with a sequencing depth of ∼50 million reads per sample.

### R-loop data analysis

CUT&Tag sequencing reads were processed using the nf-core cutandrun pipeline (v3.2.2) ^107^. Read counts were quantified for 5’ UTR regions and TPM normalized with a custom R script. Only transcripts with abundance ≥ 1 TPM in both replicates were retained for downstream analysis. Control RNAs were randomly sampled from the pool of expressed transcripts (TPM ≥ 0.1 in both replicates, excluding target genes) with the number of control RNAs matched to the number of target RNAs that passed the expression filter. RNA abundance for target and control sets was compared using log₂ counts per million (CPM) in cisplatin-treated samples.

### Cell Viability assay

A2780 cells were grown in RPMI medium with 10% FBS, and the cells were passaged at a 1:10 ratio and cultured for 72 hours before the start of the experiment. 60 µL of RPMI media containing 1 × 10^4^ cells of A2780 were seeded onto the wells of a 96-well plate and allowed to adhere for 24 hours in the incubator at 37°C. After incubation, 20 µL of the desired concentration of PhenDC3 and cPDS (0.5, 2.5, and 5 µM) were added to the adhered cells and incubated for 24 hours followed by 20 µL of serially diluted cisplatin of desired concentration (200, 100, 50, 25, 12.5, 6.25, 3.125, 1.56, 0.78, 0.39 µM) and incubated for another 24 hours. After 24 hours, 100 µL of CellTiter Glo 2.0® assay (Promega, G9242) was added to each well and incubated in the dark for 10 minutes at room temperature to stabilize the luminescent signal. The plates were then shaken on an orbital shaker for 2 minutes, and luminescence was measured using VANTAstar® microplate reader (BMG Labtech). RPMI media with 10 % FBS served as a blank. The percentage cell viability was calculated by normalizing to the wells with the lowest luminescence signal in GraphPad Prism, and IC50 values were calculated using a Nonlinear regression fit by comparing the inhibitor response to the normalized response variable slope. Each experimental condition was repeated at least 3 times. The P-values were calculated using paired Student’s t-test.

### Circular Dichroism experiments

CD samples containing 10 µM biotinylated PI4K2B (IDT, 10,392.5 Da) or PPP4R2 (IDT, 6,032.9 Da) were prepared in either annealing buffer (10 mM Tris, pH 7.5, 100 mM potassium phosphate) or annealing-prevention buffer (10 mM Tris, pH 7.5, 100 mM lithium chloride). Each preparation was brought to a final volume of 150 µl and aliquoted into 0.2 ml PCR tubes. For platination experiments, 300 µl samples at 5 µM were prepared using the same conditions and incubated with 4 mM cisplatin at 37 °C for 1 hour prior to annealing. G-quadruplex annealing was performed by heating samples to 95 °C for 5 min in a thermocycler, followed by cooling to 4 °C at a rate of 0.1 °C/s. After annealing, each 150 µL sample was combined with an equal volume (150 µL) of either 4 mM cisplatin (for platination of preformed rG4) or molecular biology–grade DEPC-treated water as a control. All mixtures were then incubated at 37 °C for 1 hour before CD measurement. CD spectra were acquired on a JASCO J-715 Circular Dichroism Spectrophotometer using a 1 mm pathlength quartz cuvette (Hellma Analytics, 100-1-40). Measurements were recorded from 320 to 200 nm with a 0.5 nm data pitch and averaged over three accumulations. Buffer-only spectra were used for background subtraction. Data averaging was performed using Microsoft Excel, and the spectra were plotted in GraphPad Prism (v10.3.1) using a 0th-order polynomial with four neighboring points on each side.

### Thioflavin T (ThT) Binding Assay

Reaction mixtures (100 µl) containing 8 µM biotinylated PI4K2B (IDT, 10,392.5 Da) or PPP4R2 (IDT, 6,032.9 Da) were prepared in either annealing buffer (10 mM Tris, pH 7.5, 100 mM potassium phosphate) or annealing-prevention buffer (10 mM Tris, pH 7.5, 100 mM lithium chloride) and dispensed into 0.2 ml PCR tubes. For platination experiments, 200 µl samples at 4 µM were prepared in the same buffers and incubated at 37 °C for 1 hour prior to annealing. G-quadruplex annealing was induced by heating to 95 °C for 5 min in a thermocycler, followed by cooling to 4 °C at a rate of 0.1 °C/s. Following annealing, each 100 µL sample was combined with 100 µL of either 1.6 mM cisplatin (post-platination) or molecular biology-grade DEPC-treated water as a control. All samples were incubated at 37 °C for 1 hour prior to mixing with ThT. The ThT working solution (2 µM) was freshly prepared from a 5 mM stock (Fisher Scientific, AC211760250) and mixed with the RNA samples in a 1:1 (v/v) ratio in a black-walled 96-well plate. Plates were shaken at 300 rpm for 5 minutes before measurement. Fluorescence was recorded in quadruplicate using a VANTAstar Microplate Reader (BMG Labtech) with excitation at 425 nm and emission scanned from 450 to 650 nm. Background subtraction was performed using buffer-only wells. Data averaging was performed using Microsoft Excel, and the spectra were plotted in GraphPad Prism (v10.3).

### Chromatin Immunoprecipitation sequencing (ChIP-seq)

A2780 ovarian cancer cells were treated with 12.5 μM cisplatin for 1 hour and crosslinked with 1% formaldehyde for 10 minutes at room temperature. Crosslinking was quenched by adding glycine to a final concentration of 0.125 M. Adherent cells were gently scraped and collected by centrifugation at 500 × g for 8 min at 4 °C. Nuclei were isolated by resuspending the cell pellet in nucleus isolation buffer (10 mM HEPES, 85 mM KCl, 1 mM EDTA, 0.5% Igepal, and protease inhibitor cocktail; ThermoFisher, PIA32965), incubating on ice for 10 minutes, and centrifuging at 500 × g for 5 min. The nuclear pellet was lysed in nuclear lysis buffer (20 mM Tris–HCl, 150 mM NaCl, 1 mM EDTA, 0.5 mM EGTA, 0.4% sodium deoxycholate, 0.1% SDS, 1% Igepal, 0.5 mM DTT, and protease inhibitors) and subjected to sonication using a Bioruptor Pico (Diagenode) for 8 cycles (30 s on, 30 s off) to shear chromatin. A 1% aliquot of the clarified lysate was reserved as an input control. Chromatin was precleared with 20 μL Dynabeads Protein G (ThermoFisher, 10003D) for 2 hours at 4 °C, then incubated overnight at 4 °C with 6 μg of anti–RNA Polymerase II antibody (clone CTD4H8; Millipore Sigma, Cat. No. 05-623-Z). Immune complexes were captured by adding 30 μL Protein G beads for 2 hours at 4 °C. Beads were sequentially washed with nuclear lysis buffer and high-salt wash buffer (10 mM Tris–HCl, 250 mM LiCl, 1% Igepal, 1% sodium deoxycholate, 1 mM EDTA, and protease inhibitors). Bound chromatin was eluted in elution buffer (10 mM Tris–HCl, pH 8.0, 1 mM EDTA, 1% SDS). The eluates were treated with RNase A (10 μg/ml) at 37 °C for 1 hour. Reverse crosslinking was performed by incubation at 65 °C overnight. The samples were finally treated with Proteinase K (20 μg/ml) at 50 ^0^C for 1 hour. DNA was purified using QIAquick Spin Columns. ChIP–seq libraries were prepared using the KAPA Hyper Prep Kit (Roche, 7962347001) following the manufacturer’s protocol, and sequenced on an Illumina NovaSeq X Plus platform (paired-end 150 bp).

### ChIP-seq data analysis

Raw sequencing reads were evaluated for quality using FastQC. High-quality reads were aligned to the GRCh38 human reference genome using Bowtie2 (v2.3.5), and alignment files were converted to BAM format, sorted, and indexed using SAMtools (v1.10). Each treatment condition (12.5 μM cisplatin or DMSO vehicle control) included two biological replicates with matched input controls. For stalled index analysis, read counts were computed using BEDTools (v2.27.1) within regions surrounding transcription start sites (TSS; −50 to +300 bp) and across gene bodies (from +300 bp to transcription end sites). Counts were normalized to reads per million per base pair (RPM/bp) using total library sizes, and background signal was corrected by subtracting input coverage.

### Western blot analysis

A2780 cells (0.5 × 10⁶) were seeded in 6-well plates and allowed to adhere for 24 hours. Cells were then treated with 12.5 µM cisplatin or DMF for 1, 3, or 6 hours. Following treatment, cells were lysed in ice-cold RIPA buffer (50 mM Tris-HCl, pH 7.4, 150 mM NaCl, 1 mM EDTA, 1% IGEPAL CA-630, 0.25% sodium deoxycholate, 0.1% SDS, and a protease inhibitor cocktail; ThermoFisher, Cat. PIA32965). Lysates were sonicated using a Bioruptor Pico (Diagenode) for 3 cycles (30 s on, 30 s off) and centrifuged at 16,000 × g for 5 min at 4 °C. The supernatant was collected, and total protein concentration was determined using the Pierce BCA Protein Assay Kit (ThermoFisher, 23227). Equal amounts of protein were mixed with 4× Bolt™ sample buffer (Invitrogen, Cat. B0007), heated at 95 °C for 5 min, and resolved on 4–12 % Bis-Tris NuPAGE™ gels (Invitrogen, NP0335BOX) at 120 V for 90 min. Proteins were transferred to 0.2 µm PVDF membranes (Bio-Rad, 1620177) at 15 V for 90 min using the Mini Blot Module (Invitrogen, B1000). Membranes were blocked for 1 hour at room temperature in 5 % non-fat dry milk in TBS containing 0.1 % Tween-20 (TBST), washed three times with TBST, and incubated overnight at 4 °C with anti-RNA polymerase II (CTD4H8; Millipore Sigma, 05-623-Z; 1:1000 in blocking buffer). β-Actin (Santa Cruz, sc-47778; 1:2000) served as a loading control. After three washes in TBST, membranes were incubated for 1 hour at room temperature with HRP-conjugated anti-mouse IgG secondary antibody (1:10,000 in blocking buffer), washed again, and developed using Pierce™ ECL Western Blotting Substrate (Thermo Fisher, 32106). Chemiluminescent signals were detected with a ChemiDoc imager (Bio-Rad Genesys Mini). Band intensities were quantified using ImageJ, normalized to β-actin, and plotted in GraphPad Prism.

### Total RNA sequencing

A2780 ovarian cancer cells were treated with 10 μM cisplatin or vehicle (DMF) for 24 h, in three independent biological replicates per condition. Total RNA was isolated using the Zymo Quick RNA MiniPrep Kit (Zymo Research, R1054) with on-column DNase I digestion, following the manufacturer’s protocols. RNA was quantified by NanoDrop. RNA-seq libraries were prepared from 1 µg of total RNA after the removal of rRNA, using the KAPA RNA HyperPrep Kit with HMR RiboErase (Roche, KR1351 v4.21), and sequenced on an Illumina NovaSeq X Plus platform to generate ∼50 million paired-end reads. Raw sequencing data were quality-checked using FastQC, and adapter sequences were trimmed using Cutadapt (minimum read length = 1 bp). Trimmed reads were aligned to the human reference genome (GRCh38, GENCODE v46 annotation) using STAR with local alignment parameters allowing up to 100 multi-mapped reads. Gene and isoform abundance were quantified with RSEM (v1.3.1) ^108^. Across all samples, 18,843 transcripts were detected at TPM > 1 in at least two of the three cisplatin-treated replicates, corresponding to 11,289 unique genes.

## Conflict of Interest

The authors declare no conflict of interest.

## Funding

This work was supported by the Department of Defense [OC220133 to S.J.N. and V.J.D.]; National Science Foundation [CHE2109255 to V.J.D.]; and National Institutes of Health [R35GM150636 to S.J.N.]. The Biacore Molecular Interaction Shared Resource (BMISR) facility at Georgetown University is supported by the National Institutes of Health [P30CA51008 and 1S10OD019982-01]. ICP-MS measurements were performed in the OHSU Elemental Analysis Core with partial support from the National Institutes of Health [S10OD028492].

## Supplementary Data

**Supplementary Figure 1.**
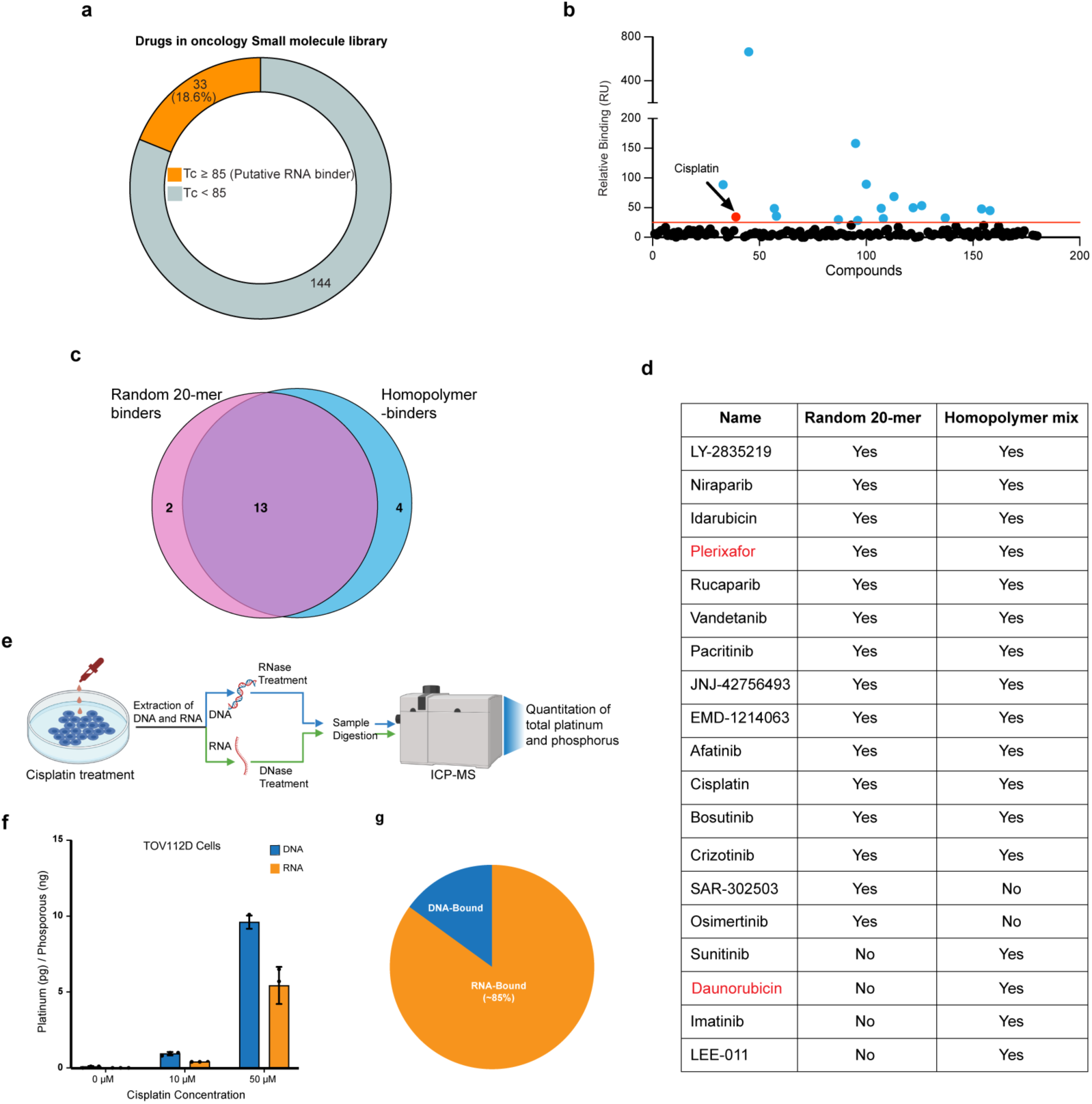
Screening of RNA binding oncology small molecules. (a) Chart showing the fraction of predicted RNA-binders in the anticancer small molecule library (NCI/DTP AOD XI) used in the screening experiment. (b) The binding of individual small molecules in the library to the equimolar 12-mer homopolymer RNA pool. The red line is the 25% threshold used as a cut-off. (c) Overlap hits between random-20mer RNA ligands and equimolar concentration of homopolymers. (d) Table describing the hits obtained by independent screening using random 20-mer RNAs and equimolar homopolymer-mix. Small molecules included in the combined list of known RNA-binding small molecules (Supplementary Table 1) are highlighted in red. (e) Cartoon depicting nucleic acid fractionation and ICP-MS. (f) Quantitation of cisplatin accumulation in the DNA and RNA fractions of TOV112D cells. (g) Relative distribution of cisplatin in cellular RNA and DNA fractions, extrapolated from ICP-MS and RNA and DNA content in mammalian cells (29,30).

**Supplementary Figure 2.**
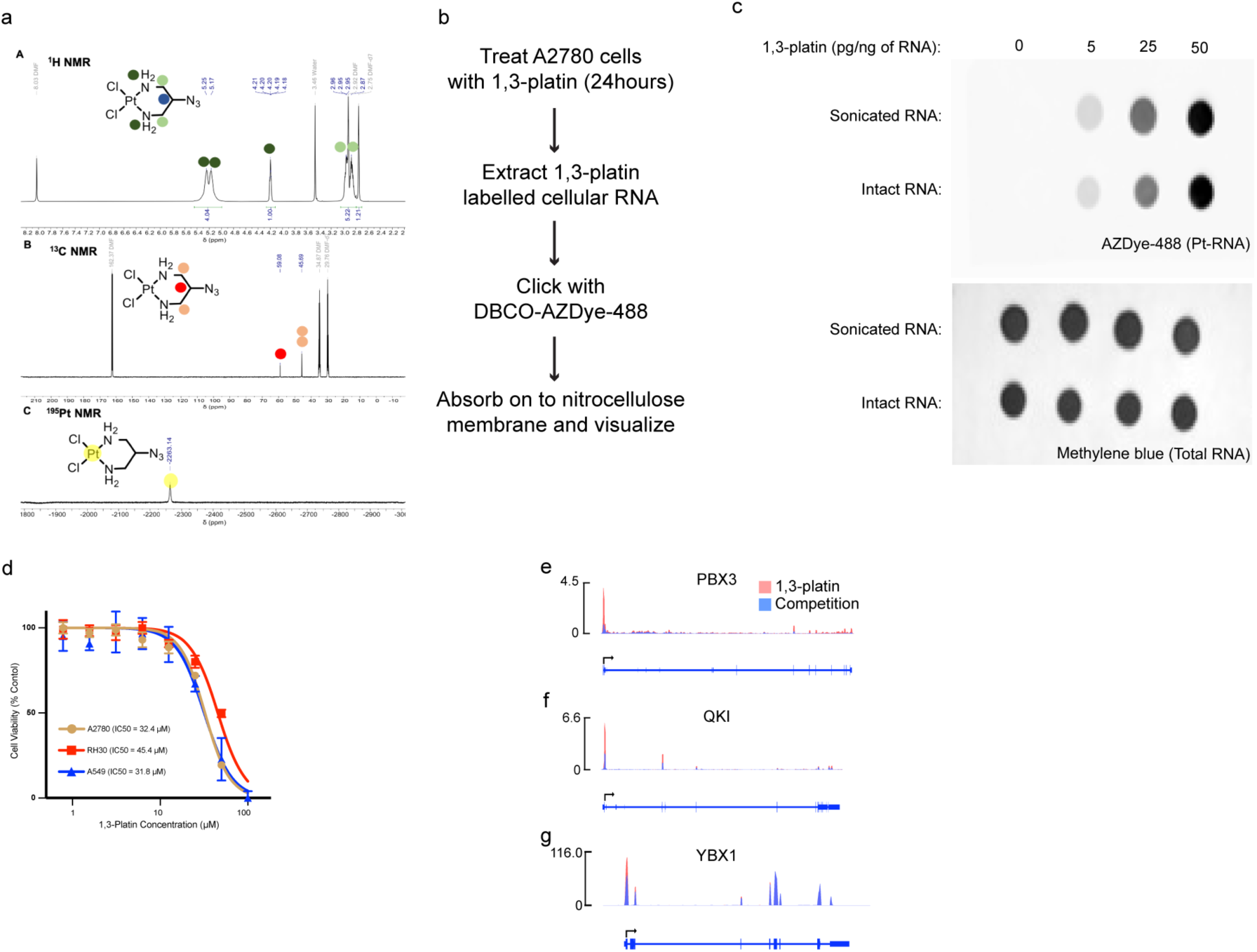
Development of Plat-RNA seq. (a) NMR data showing the purity of 1,3-platin. (b) Outline of in vitro click-labelling experiment. (c) Dot-blot showing efficient click-reaction to 1,3-platin labelled RNAs with DBCO-AZDye-488 and an increase in signal with increased RNA labeling. Methylene blue labelling of the membrane shows equal loading of the RNAs. (d) Cell viability assay to estimate the potency (IC50) of 1,3-platin in the indicated cell lines. (e), (f), (g) Read density tracks along the transcripts for the genes PBX3 (e), QKI (f), and YBX1 (g) in A2780 cells.

**Supplementary Figure 3.**
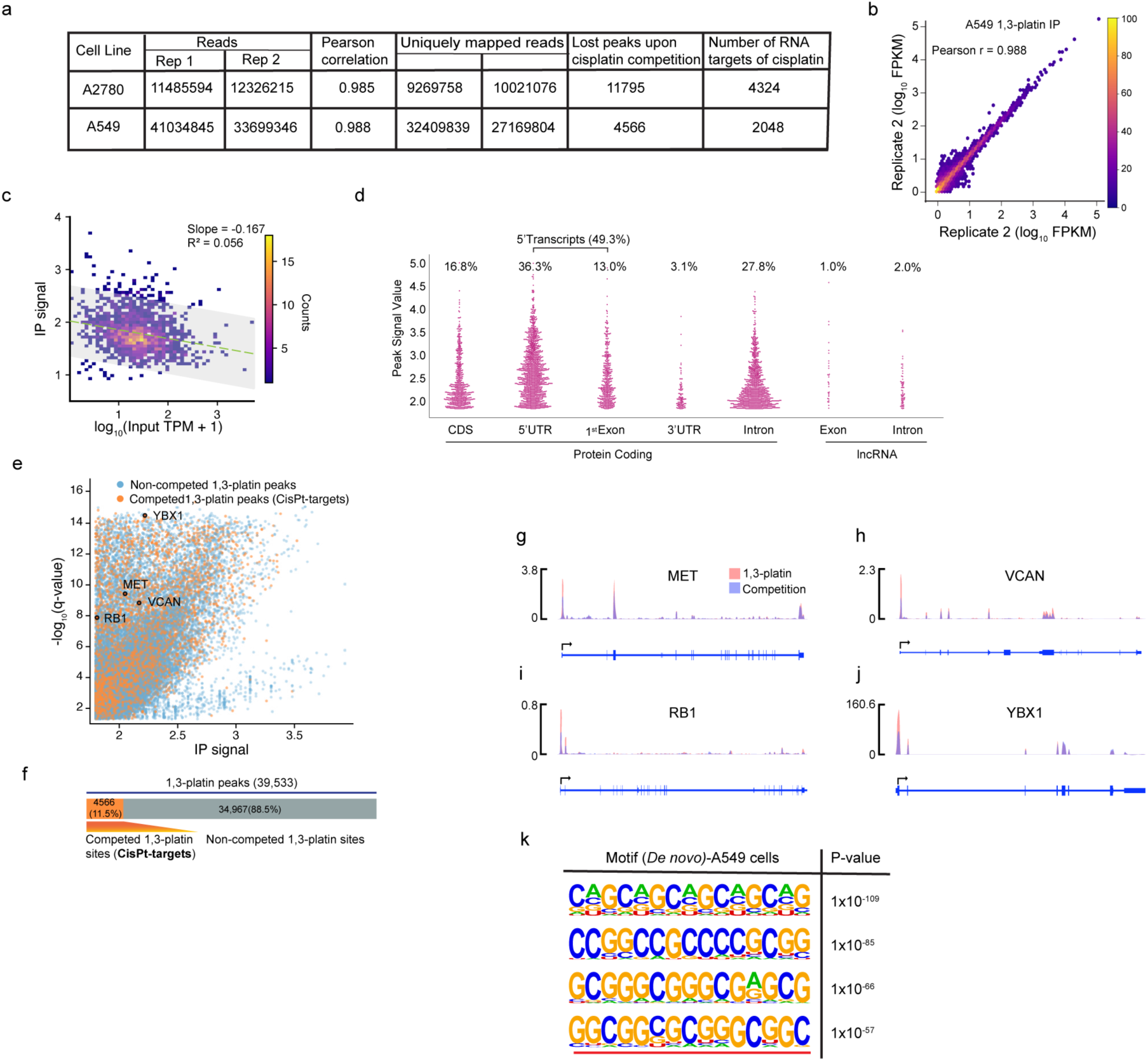
Plat-RNAseq analysis of A549 cells. (a) Table showing Plat-RNAseq sequencing quality information in the indicated cell lines. (b) Scatter plot of 1,3-platin IP replicates reads signal from A549 cells. The r-value was derived from the Pearson correlation analysis. (c) Scatter plot showing Plat-RNAseq pull-down signals versus RNA expression levels in A549 cells, revealing no correlation between transcript abundance and pull-down efficiency (R2 = 0.056). (d) The relative distribution of 1,3-platin binding across different transcript locations in A549 cells. (e) Plat-RNAseq data visualized as dot plots. Each dot represents a 1,3-platin-interacting RNA fragment identified through peak calling. RNA fragments showing reduced binding (Log2FC< -0.5) are designated as cisplatin targets (CisPt-targets) and highlighted in orange. Four CisPt-targets, for which read density plots are shown, are circled. (f) Quantitation of 1,3-platin-RNA peaks showing the number of sites that were either competed or not competed by cisplatin. (g), (h), (i), (j) Read density tracks along the transcripts for the genes MET (g), VCAN (h), RB1 (i), and YBX1 (j) in A549 cells. (k) Motif analysis of cisplatin-enriched sequences in A549 cells reveals G-rich motifs. The motif underlined in red is a predicted rG4.

**Supplementary Figure 4.**
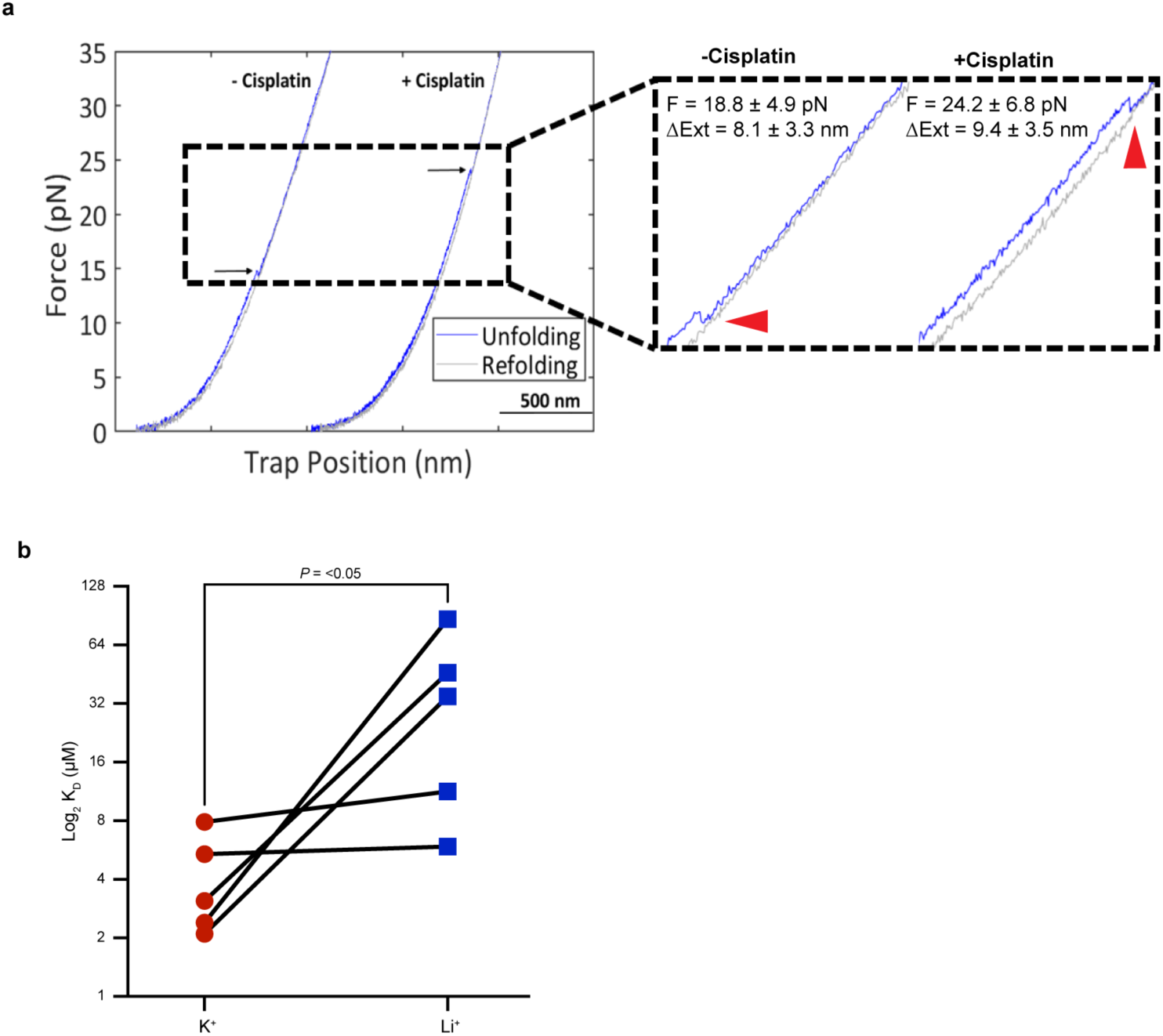
Cisplatin exhibits higher affinity binding to rG4 RNA and modulates its structural stability. (a) Zoomed-in optical tweezer force-extension curve of PI4K2B rG4 in the absence and presence of cisplatin. The arrows point to the major unfolding events in these two trajectories. (b) Spaghetti plot showing paired binding affinity measurements of the PI4K2B RNA oligo under rG4-stabilizing (K+) and rG4-disrupting (Li+) buffer conditions (n = 5). P-value was calculated using a paired Student’s t-test.

**Supplementary Figure 5.**
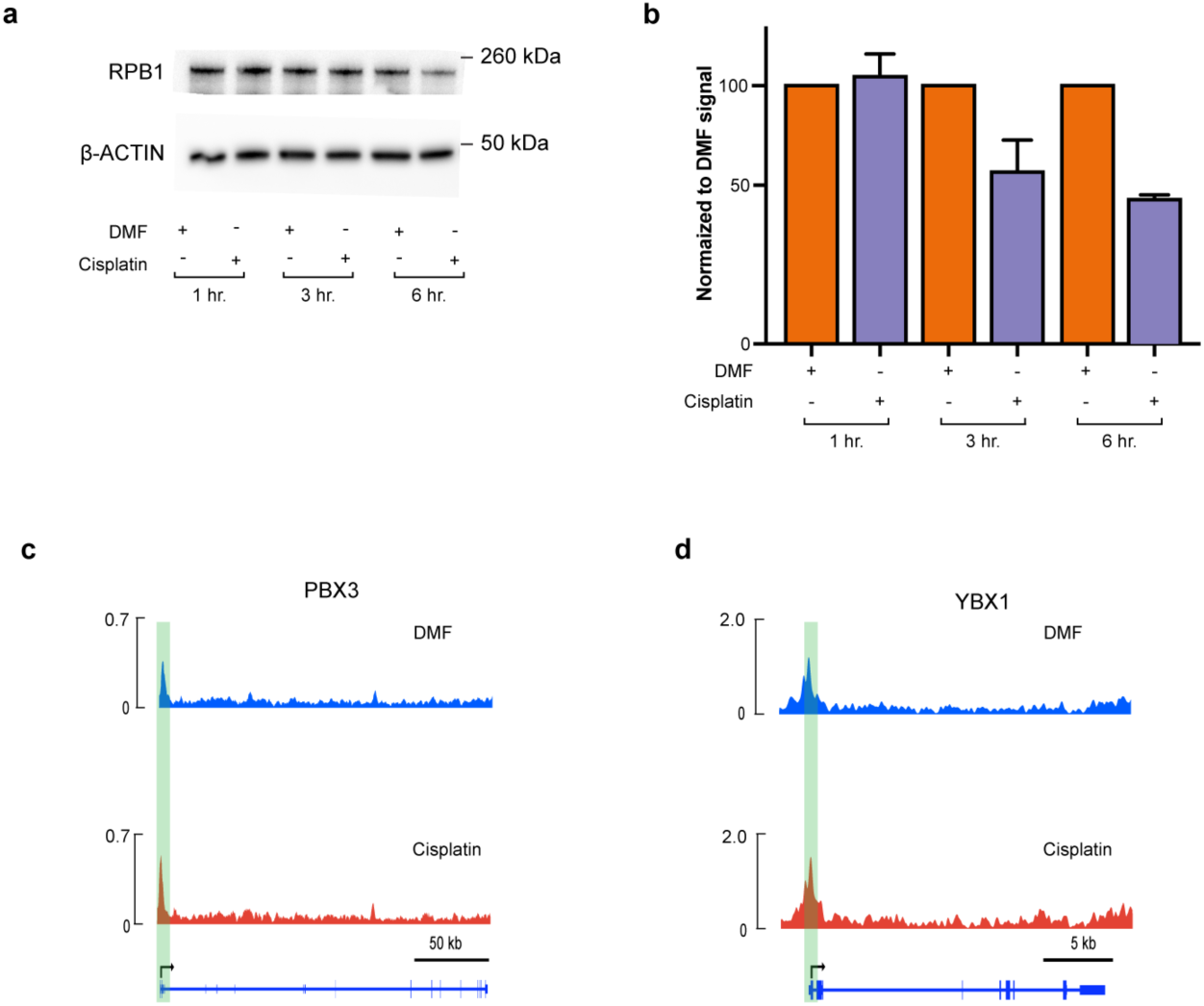
Cisplatin effect on RNA Pol II stability and stalling. (a) RNA Pol II Western blot showing degradation of RNA Pol II after 3 hours and 6 hours of cisplatin treatment. Pol II level remains unchanged after 1 hour of treatment, hence used as a timepoint for ChIP-seq experiments. (b) Quantitation of the Western blot data. (c), (d) Read density tracks of RNA Pol II ChIP-seq at PBX3 (c) and YBX1 (d) loci. The promoter proximal region is highlighted in green.

**Supplementary Figure 6.**
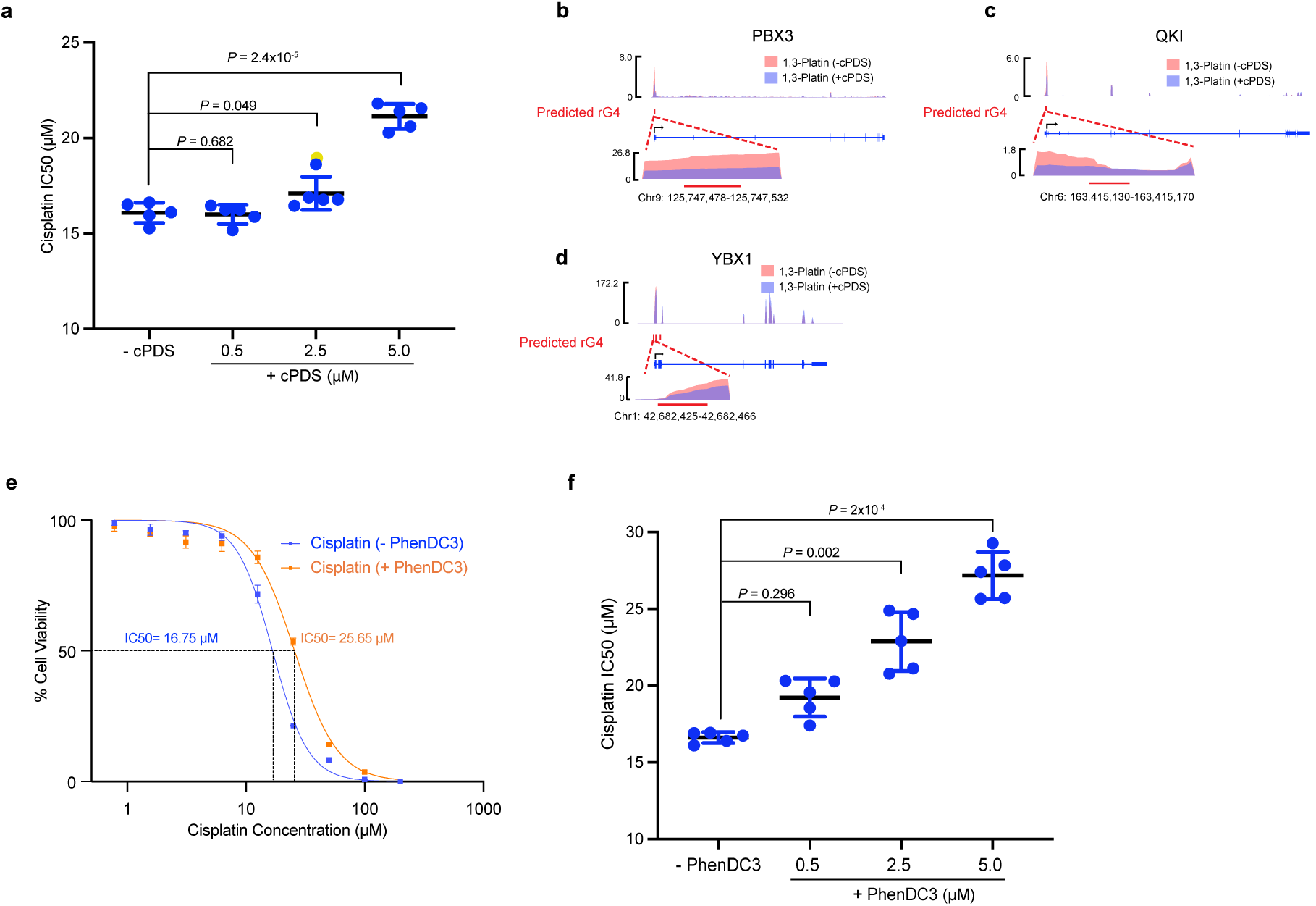
Cisplatin-rG4 accumulation promotes cellular cytotoxicity. (a) Distribution of cisplatin IC50 change, samples treated with vehicle (-cPDS) and pre-treated with indicated concentration of cPDS in replicate experiments. P-value was calculated using Student’s t-test. (b), (c), (d) Read density tracks along the transcripts for the genes, PBX3 (b), QKI (c), and YBX1 (d) in A2780 cells from 1,3-platin-pulldown sequencing experiments. The cells were treated with vehicle (-cPDS, pink) or pretreated with cPDS (+cPDS, purple) before treatment with 1,3-platin. The rG4 sites are zoomed-in. (e) Cell viability assay showing reduced cisplatin efficacy (increased IC50) in cells pretreated with the rG4-ligand PhenDC3. (f) Distribution of cisplatin IC50 change, samples treated with vehicle (-PhenDC3) and pre-treated with indicated concentration of PhenDC3 in replicate experiments. P-value was calculated using Student’s t-test.

## Notes

### Competing Interest Statement

The authors have declared no competing interest.

### Summary of Updates

Figure 6 from the first version was removed to align with the manuscript's current focus on demonstrating the contribution of the cisplatin-RNA interaction to its cytotoxic mechanism.

